# A nanobody-based degron system for targeted protein knockdown in *Dictyostelium discoideum*

**DOI:** 10.1101/2025.10.23.684119

**Authors:** Hidenori Hashimura, Shoko Fujishiro, Nao Shimada, Tomoko Adachi, Toyoko Sugita, Satoshi Kuwana, Satoshi Sawai

## Abstract

**Backgrounds:** The cellular slime mold *Dictyostelium discoideum* is a widely used model system for studying basic processes in cell and developmental biology. While genetic tools, such as targeted gene disruption by homologous recombination and genome editing using CRISPR/Cas9, are well-established in *D. discoideum*, efficient methods for conditional loss-of-function studies are limited. Here, we developed a nanobody-based degron system for *D. discoideum* based on ALFA-tagged protein recruitment to the Skp1-Cullin-F-box (SCF) complex.

**Results:** ALFA-tagged Histone H1 was efficiently degraded by expressing anti-ALFA nanobody (NbALFA) fused to the *D. discoideum* FbxD F-box domain (‘dictyGrad-ALFA’). Cell type-specific targeting was achieved by expressing dictyGrad-ALFA under prestalk- and prespore-specific gene promoters. Furthermore, targeting of adenylyl cyclase A (ACA) resulted in the expected aggregation-deficient phenotype, validating the efficacy of dictyGrad-ALFA-mediated protein depletion. Cell type-specific ACA degradation delayed development but eventually resulted in normal fruiting bodies. Our ALFA-tag approach was further used for conditional knockdown in combination with the auxin-inducible degron 2 (AID2) system, which relies on indole-3-acetic acid (IAA)-dependent binding between NbALFA-mAID and a OsTIR-F-box-Skp1A fusion protein. We obtained efficient IAA-induced degradation in prestalk cells; however, efficiency was low in other cell types.

**Conclusions:** Together, these systems pave the way for conditional and cell type-specific protein degradation in *D. discoideum*, enabling functional analyses of essential genes for development and survival.

## Background

The cellular slime mold *Dictyostelium discoideum* is a unique amoebozoan species that is highly amenable to molecular genetic approaches in the fields of cell and developmental biology and evolution. Its haploidy and ease of cell culture facilitate efficient gene knockout, allowing in-depth molecular and genetic analyses of basic cellular and developmental processes. Gene knockout in *D. discoideum* has traditionally been achieved by inserting a drug resistance cassette in the gene of interest by homologous recombination [1–3]. More recently, CRISPR/Cas9-mediated genome editing has been utilized for loss-of-function studies [4, 5]. In combination with random mutagenesis screens, knockout studies in *D. discoideum* have successfully uncovered genes including, but not limited to, those important for cytokinesis, cell migration, cell-cell signaling, cell differentiation, morphogenesis [6–10]. However, simple gene-knockout strategies have various limitations. For example, they cannot be applied to genes essential for the growth phase of the organism. Similarly, the loss of genes that are essential for early development precludes comprehensive analyses of their roles in later developmental stages. Moreover, the generation of knockout cells may have secondary effects, such as altered cell homeostasis and changes in gene expression profiles [11–13], potentially confounding the interpretation of the mutant phenotype. Posttranscriptional gene silencing through RNA interference is not widely used in *D. discoideum,* in part because RNAi is under developmental control [14]. There are also general limitations stemming from off-target gene silencing, which affects non-target transcripts that share only partial sequence complementarity with the siRNA [15, 16]. Efficient knock-down methods for *D. discoideum* are therefore needed.

Recent advances in nanobody-based degron systems [17] offer a promising approach for targeted protein degradation and knockdown studies in *D. discoideum*. These methods exploit the highly conserved ubiquitin–proteasome pathway in which the SCF (Skp1-Cullin-F-box protein) complex mediates the key ubiquitination step that brings together E3 ubiquitin ligases and the protein to be degraded [18]. By replacing the WD40 repeat in an F-box protein with a nanobody against GFP, a protein of interest (POI) tagged with GFP can be directed toward ubiquitination by the SCF complex, thus facilitating stage- and cell-type-specific loss-of-function studies [19, 20]. In the fruit fly (*Drosophila melanogaster*) ‘deGrad’ system [19], Gal4/UAS-based expression of an anti-GFP nanobody (vhhGFP4) fused to the F-box domain of Slmb targets GFP-tagged proteins for degradation. This has been successfully demonstrated for a GFP-tagged myosin regulatory light chain, the transcription factor Ap, and the transmembrane protein Crb at specific stages of embryonic development. A similar ‘zGrad’ system [20] in zebrafish (*Danio rerio*), which employs the F-box domain of its native Fbxw11b, induces the degradation of GFP-tagged Cadherin, α-E-Catenin, and the chemokine receptor Cxcr4b during embryogenesis.

The strategies described above are slow-acting, with degradation often taking several hours, and are difficult to terminate once initiated. For more responsive targeted protein degradation, proteolysis targeting chimeras (PROTACs) employ chimeric molecules composed of a compound that directly binds a POI and a ligand that recruits an SCF complex component [21]. Similarly, the dTag system targets a POI tagged with FKBP(F36V) using a chimeric cell-permeable compound that bridges a substrate receptor of an SCF complex component (CRBN) and FKBP(F36V) (e.g., dTag-13) [22]. These approaches require a compound designed to bind to specific components of the SCF complex, which can be context- and species-dependent. Alternatively, the auxin-inducible degron (AID) system uses indole-3-acetic acid (IAA)-dependent binding between a POI fused to mAID tag and the *Oryza sativa* TIR protein (OsTIR) [23]. Depending on the timing and location of TIR-F-box domain fusion protein expression, a POI can be degraded in a time- and IAA dose-dependent manner. A more recent AID2 system utilizes a modified OsTIR that creates a cavity in its auxin-binding pocket, which in combination with the ligand IAA analog 5-Ph-IAA improves specificity and reduces background degradation [24]. AID2 has been implemented in various organisms, including yeast, nematode, fruit fly, and mouse [24–27].

In this study, we developed dictyGrad-ALFA, a nanobody-based degron system for targeted protein degradation in *D. discoideum.* The system targets a POI tagged with a synthetic ALFA-tag and a fusion protein comprising an anti-ALFA-tag nanobody and the *D. discoideum* FbxD F-box domain. We demonstrate efficient and cell type-specific degradation of ALFA-tag-fused Histone H1-mScarlet-I in both vegetative and multicellular stages. The efficacy of the approach is further demonstrated by knocking down adenylyl cyclase A (ACA). Furthermore, we extended the approach to the AID2 system for inducible protein depletion.

## Materials & Methods

### Plasmid construction

The primers and plasmids used in this study are listed in Supplementary Tables 1 and 2. Phusion polymerase (New England Biolabs, Ipswich, MA, USA) and KOD -Plus- Neo (TOYOBO, Osaka, Japan) were used for PCR amplification. PCR-amplified DNA was fused to linearized vectors using the In-Fusion Snap Assembly Cloning Kit (TAKARA, Kusatsu, Japan). T4 DNA ligase (Ligation High Ver. 2; TOYOBO) was used for general DNA ligation. For constitutive expression of fluorescent protein-tagged NbALFA, mAID, OsTIR, and extrachromosomal plasmids with the *coaA* and *act15* promoters were used. The *D. discoideum* codon-optimized DNA sequences for ALFA-tag [28], NbALFA [28], mAID [29], and OsTIR [23] are summarized in the Supplementary materials.

To obtain vectors for nanobody expression, #HH574, #HH623 and #HH686, NbALFA was amplified and inserted at BglII or SpeI sites of the following plasmids: #HH242 pDM358_*act15p*:mScarlet-I-(GGS)_2_-MCS (for #574), #HH162 pDM326_*act15p*:Achilles-(GGS)_2_-MCS [30] (for #HH623), and #HH675 pDM304_*act15p*:Electra2-(GGS)_2_-MCS [31] (for #HH686). To obtain vectors that drive the expression of Achilles-NbALFA (#HH707) and Electra2-NbALFA (#HH686) under the *coaA* promoter, PH_Akt_-mNeonGreen fragments flanked by BglII and HindIII sites in the #HH109 pDM304_*coaAp*:PH_Akt_-mNeonGreen [30] were replaced with Achilles-NbALFA-*act8*t or Electra2-NbALFA-*act8*t sequences. To obtain an expression vector for NbALFA fused to the mAID tag (#HH676), Dictyostelium-codon-optimized DNA for mAID was fused to the N-terminus of NbALFA by overlap PCR extension. The amplified sequence of mAID-NbALFA was cloned into pDM304 [32] at BglII and SpeI sites.

A partial coding sequence of FbxD (FbxDΔWD, 1–198 aa) including the F-box domain was amplified from genomic DNA and inserted into the BglII site of #HH707 to obtain plasmid #HH708. The *ecmAO* and *D19* promoters were cloned from genomic DNA [30] and inserted at the XhoI and BglII sites of pDM304. To construct vectors for the expression of Achilles-FbxDΔWD-NbALFA (dictyGrad-ALFA) under the *ecmAO* (#HH657) or *D19* promoter (#HH659), respectively, the sequence Achilles-FbxDΔWD-NbALFA-*act8t* was amplified and inserted at the BglII and HindIII sites.

To generate the knock-in vector #HH608, the Histone H1-mScarlet-I-ALFA fragment was PCR-amplified with primers containing the ALFA-tag sequence. The purified product was inserted at the BglII and SpeI sites of pDM1501 [33]. To construct the CRISPR/Cas9 vector #HH681, a gRNA oligo targeted to the *acaA* locus (4515–4532 bp from the start codon, Supplementary Table 2) was annealed and inserted into pTM1285 [4] using Golden Gate Assembly with BpiI (Thermo Scientific, Waltham, MA, USA) and T4 DNA ligase (New England Biolabs). Cas-Designer [34] was used to design sgRNA sequences.

To construct OsTIR(F74G) expression vectors, the F74G mutation was introduced in the codon-optimized OsTIR DNA by overlap extension PCR. The amplified fragment was sub-cloned in the pCR Blunt II-TOPO vector (Zero Blunt TOPO PCR Cloning Kit; Invitrogen, Waltham, MA, USA). The mutation was confirmed by a sequence analysis. To obtain the OsTIR(F74G) expression vector (#HH697), the PH_Akt_ fragment in pDM304_*coaAp*:PH_Akt_-mNeonGreen (#HH109) [30] was replaced with the OsTIR(F74G) fragment at the BglII and SpeI sites, and mNeonGreen was replaced with Achilles at the SpeI and HindIII sites. To obtain a vector carrying the Skp1A sequence (#HH350), the promoter and coding sequence of *Skp1A* were amplified from genomic DNA and inserted into pDM358_ *act15p*:MCS-(GGS)_2_-mScarlet-I (#HH253) [30] at the SpeI and XhoI sites, removing the *act15p* promoter. To obtain the Skp1A-(GGS)_2_-OsTIR(F74G) expression vector (#HH698), PCR-amplified fragments of OsTIR(F74G) with the (GGS)_2_ linker were inserted at the SpeI and HindIII sites of vector #HH350, replacing the mScarlet-I-*act8term* fragment. To generate an expression vector for Skp1A-(GGS)_2_-OsTIR(F74G) (#HH698), the Skp1A-(GGS)_2_-OsTIR(F74G) fragment was amplified and inserted into vector #HH697 at the BglII and SpeI sites, thereby placing the OsTIR(F74G) fragment. To obtain dual expression vectors for OsTIR(F74G) or Skp1A-(GGS)_2_-OsTIR(F74G) and mAID-NbALFA expression (#HH699 and #HH700), a fragment containing the *act15* promoter, the coding sequence of mAID-NbALFA, and the *act8* terminator region were PCR-amplified from vector #HH676 and inserted at the NgoMIV site of #HH697 and #HH698, respectively.

To obtain the mTagBFP2 C-terminus-ALFA tag vector (#HH570), the mTagBFP sequence was PCR-amplified to include the ALFA-tag sequence at the 3′ end and inserted in the SpeI site of vector pDM304. To obtain Histone H1-mtagBFP2-ALFA and ACA-mScarlet-I expression vectors (#HH573 and #HH722), the coding sequences of Histone H1 and ACA were PCR-amplified using genomic DNA as a template and inserted into the BglII and SpeI cloning sites of vector #HH570 and vector pDM358_*act15p*:MCS-(GGS)_2_-mScarlet-I (#HH253) [30], respectively.

### Transformation

*Dictyostelium discoideum* Ax4 was used as a parental strain. To introduce extrachromosomal vectors, cells were electroporated with 1 µg of a vector following the standard protocol [35]. For the knock-in of Histone H1-mScarlet-I-ALFA at the *act5* locus, a 4.6 kbp fragment that included the *act5* homology arms, fluorescence protein-coding sequence, and hygromycin-resistant cassette was excised from #HH608 with NgoMIV. Cells were electroporated together with 10 µg of the purified fragment. For *acaA* knockout, cells were electroplated with 5 µg of the vector pTM1224 [4]. To generate a knock-in of mScarlet-I-ALFA at the *acaA* locus via CRISPR/Cas9, cells were electroporated with 10 µg of the vector #HH681 and 2 µg of PCR-amplified fragments of mScarlet-I-ALFA flanked by 60 bp homology arms (Supplementary Table 2) as a donor template following a previously described protocol [36].

Transformants were selected in HL5 medium containing 10 μg/mL G418 or 60 μg/mL Hygromycin B either alone or in combination. To isolate a single clone, approximately 50–70 cells were plated on the half-diluted SM agar plate (Dextrose, Peptone, Yeast Extract, MgSO_4_, KH_2_PO_4_, Na_2_HPO_4_ [37]) with *Klebsiella aerogenes* and incubated at 22 °C until plaques formed. To verify mutations introduced by CRISPR/Cas9 genome editing, the target sequence was amplified by PCR using cell lysates as a template and sequenced. Cell lysates were prepared by picking cells from isolated plaques suspended in lysis buffer (10 mM Tris pH8.5, 50 mM KCl, MgCl_2_, 0.45% NP40, 0.45% Tween20 [38]) containing Proteinase K (QIAGEN, Netherlands) and heated at 56 °C for 10 minutes and at 95 °C for another 10 minutes. To obtain cell lines expressing Achilles-NbALFA, Achilles-FbxD Δ WD-NbALFA (dictyGrad-ALFA), or Skp1A-(GGS)2-OsTIR(F74G)-Achilles at a high level, colonies picked from SM agar plates were checked for fluorescence levels using a confocal microscope. Isolates with high fluorescence signals were chosen and maintained in HL5 medium containing 40 μg/mL G418.

### Cell culture, development, and fixation

Cells were grown in HL5 medium at 22 ℃ either as a suspension culture in shaken flasks or as adherent culture in Petri dishes. The growth medium contained 10 or 40 µg/mL G418 and 60 µg/mL Hygromycin B where appropriate. To image the vegetative stage, cells were collected, washed twice, and resuspended in phosphate buffer (PB: 12 mM KH_2_PO_4_, 8 mM Na_2_HPO_4_, pH 6.5). An aliquot of the cell suspension was plated directly onto a 𝜙25 mm round coverslip (Matsunami, Osaka, Japan) mounted on a metal chamber (Attofluor, Invitrogen, Carlsbad, CA, USA). The cells were allowed to attach to the substrate before image acquisition. To obtain images of slugs and fruiting bodies, cells were starved and allowed to develop as described previously [30]. In brief, washed cells were suspended at a density of 2 × 10^7^ cells/mL in PB, and 5 µL of the cell suspension was deposited on an agar plate (2% agar (Bacto) in Milli-Q water). After 15–20 hours, samples were excised together with the agar substrate and placed upside-down on a coverslip with a polyethylene terephthalate spacer ring with a 50 µm height (vinyl patch transparent Ta-3N; Kokuyo, Osaka, Japan). To label cellulose (Fig. 4B and 5C), cells were developed on agar containing 0.1 mg/mL Fluorescent Brightener 28 (Calcofluor white; MP Biomedicals, Irvine, CA, USA, 158067) [30].

A 10 mM 5-Ph-IAA (MedChemExpress, Monmouth Junction, NJ, USA) stock solution was prepared in DMSO. For auxin-induced protein degradation in vegetative cells, 5-Ph-IAA stock solution was diluted 1:1000 or more in 1 mL of HL5 medium. An equivalent final volume of DMSO was used as a mock control. For the IAA-induced degradation of ACA-mScarlet-I-ALFA in starved cells (Supplementary Fig. 7), growing ACA-mScarlet-I-ALFA knock-in cells were washed and suspended in 1 mL of Developmental buffer (DB: 6 mM KH_2_PO_4_, 4 mM Na_2_HPO_4_, 2 mM MgSO_4_, 0.2 mM CaCl_2_, pH 6.5) at a density of 10^6^ cells/mL. Then, 1 µL of 5-Ph-IAA stock solution or DMSO was added. An aliquot of the cell suspension was plated on a round coverslip mounted on a metal chamber. The sample was incubated for 24 hours before observations.

For Hoechst 33342 staining, 1 mL of the cell suspension in HL5 medium at a density of approximately 4 × 10^5^ cells/mL was deposited on a round coverslip mounted on a metal chamber. Cells were allowed to settle and attach to the glass surface for 10 minutes. After removing the medium, 3.7% formaldehyde in phosphate-buffered saline (PBS: 137 mM NaCl, 2.7 mM KCl, 10 mM Na_2_HPO_4_, 1.8 mM KH_2_PO_4_, pH7.4) was added to the chamber and incubated for 10 minutes at 22 ℃. For the multicellular stage, washed cells were suspended at a density of 1 ×10^7^ cells/mL in PBS, and 160 µL of the cell suspension was deposited on a 1.2 × 1.2 mm area of a mixed cellulose ester filter (A045P047A; ADVANTEC TOYO, Tokyo, Japan). The filter was immediately mounted on a 𝜙 37 mm paper pad (M-085; ADVANTEC TOYO) soaked with 1 mL of Milli-Q water containing 10 µM 5-Ph-IAA. After 20 hours, the filter with developing slugs was fixed whole-mount by soaking it in 3.7% formaldehyde in PBS for 15 minutes at approximately 23 °C. The samples were washed with PBS three times and stained with 0.4 µg/mL Hoechst 33342 (Dojindo Laboratories, Kumamoto, Japan) in PBS for 30 minutes at 22 ℃. The sample was washed with 500 µL of PBS three times.

### Image acquisition and analysis

All images were acquired at 22 ℃. For fluorescence images, an inverted microscope (IX83; EVIDENT, Tokyo, Japan) equipped with a multibeam confocal scanning unit (CSU-W1; Yokogawa, Tokyo, Japan) and CMOS cameras (ORCA-Fusion BT; Hamamatsu Photonics, Hamamatsu, Japan) was employed. 405, 488 and 561 nm lasers were used for excitation, together with a multiband dichroic mirror (T405/488/561/640 nm) and 447/60, 525/50 and 617/73 bandpass filters for detection. For fluorescence imaging in Supplementary Fig 1, an inverted microscope (IX83; Evident) with a confocal scanning unit (CSU-W1; Yokogawa) and an electron-magnified CCD camera (iXon 888; Andor, Belfast, Northern Ireland, UK) were used with 405 and 561 nm lasers with appropriate dichroic beam splitters and emission filters, as described previously [30]. Images of slug samples were acquired using a stereomicroscope (SZX16; EVIDENT) equipped with a CMOS camera (DP23-CU-1-2; EVIDENT) and a sharp-cut filter (R60-40; CCS, Kyoto, Japan). A stereomicroscope (SZX10; EVIDENT) with a digital camera (EOS 700D; Canon, Tokyo, Japan) was also used.

Images were analyzed using ImageJ (Fiji distribution 2.16.0) and R 4.0.2 statistical packages. The fluorescence intensity of Histone H1-mScarlet-I in the nuclear regions was extracted from nuclear masks. The masks were generated based on Hoechst 33342 staining with Trainable Weka Segmentation [39]. In all statistical analyses, *P-*values were determined using the Wilcoxon rank-sum test with Bonferroni adjustment. The dose-response depletion and recovery in Fig. 6E and G were fitted with non-linear regression using four parameters. The time-course data in Fig. 6F were fitted to an exponential decay function.

### Western Blotting

For western blotting, 10 µL of the cell suspension (1 × 10^8^ cells/mL) was added to 10 µL of lysis buffer (20 mM HEPES, 200 mM KCl, 1% Triton-X, 0.4% NP-40, 0.03% β-mercaptoethanol, 1x complete Protease Inhibitor Cocktail). Then, 20 µL of the lysate was mixed with 10 µL of SDS sample buffer and boiled at 95 ℃ for 5 minutes. For each sample, 20 µL was loaded on a 4–20% polyacrylamide gel (Mini-PROTEAN TGX; BioRad, Hercules, CA, USA) and blotted onto a PVDF Membrane (Amersham Hybond P 0.45μm; Cytiva, Marlborough, MA, USA). The membrane was blocked with 1% skim milk in TBST and probed with an Anti-RFP mouse monoclonal antibody (6g6; Proteintech, Rosemont, IL, USA) at a 1:1000 dilution as the primary antibody followed by a fluorescent anti-mouse-IgG goat antibody (StarBrightBlue700; BioRad) at a 1:2500 dilution as the secondary antibody. The membranes were later re-probed with an anti-actin mouse monoclonal antibody (MA5-11869; Invitrogen) at a 1:250 dilution. A chemiluminescence imager (iBright1500; Invitrogen) was used to acquire gel and blot images.

## Results

### Nanobody-based targeted protein degradation in *D. discoideum*

To establish a targeted protein degradation system in *D. discoideum*, we adopted a nanobody-based approach similar to those deployed successfully in the fruit fly *D. melanogaster* (deGradFP) *[19]* and zebrafish *D. rerio* (zGrad) [20]. These systems employ an anti-GFP nanobody (vhhGFP4) [40] fused to the F-box domain of species-specific F-box protein Slmb (for fruit fly) *[19]* and Fbxw11b (for zebrafish) [20] to target the degradation of GFP-tagged proteins. Due to relatively large size, however, GFP-tagging can affect the function of the POI. Moreover, in some cases, it is desirable to reserve GFP for imaging purposes. We therefore used ALFA-tag and its nanobody, NbALFA [41, 42]. ALFA-tag is a small (15 aa) hydrophilic, uncharged artificial alpha-helix designed to serve as a unique epitope tag in prokaryotic and eukaryotic host systems [28]. In *D. discoideum,* the ALFA-tag has been used to purify and study the lysozyme-like protein AlyA [43]. First, to verify the ALFA-tag/NbALFA association in vivo, we constructed *D. discoideum* strains that co-express mScarlet-I-NbALFA and Histone H1-mTagBFP2 with or without the ALFA-tag. Both Histone H1-mTagBFP2 and Histone H1-mTagBFP2-ALFA were localized to the nucleus (Supplementary Fig. 1) [30]. In cells expressing Histone H1-mTagBFP2, mScarlet-I-NbALFA was distributed in both the cytosol and nuclei (Supplementary Fig. 1; upper panel). mScarlet-I-NbALFA was confined mostly to the nuclei in Histone-mTagBFP2-ALFA-expressing cells (Supplementary Fig. 1; lower panel). These results indicate that *D. discoideum* nucleoplasm supports the association between NbALFA and ALFA-tag.

We next evaluated the nanobody fused to the F-box domain and its ability to induce the degradation of an ALFA-tagged protein. A knock-in strain carrying Histone H1-mScarlet-I-ALFA at the *act5* locus was generated as a tester. In this strain, we introduced a plasmid for the constitutive expression of ‘dictyGrad-ALFA’ (Fig. 1A, B), NbALFA fused to the N-terminus of FbxD. FbxD is a *D. discoideum* F-box protein (Fig. 1A) known to directly interact with Skp1A and Skp1B [44]. In the growth phase, the fluorescence signal of Histone H1-mScarlet-I-ALFA in cells expressing dictyGrad-ALFA was 73% lower than that in cells without nanobody expression (Fig. 1C, D). Consistent with these findings, western blotting against mScarlet-I showed that the cells expressing dictyGrad-ALFA exhibit a 77% reduction in Histone H1-mScarlet-I-ALFA protein (Fig. 1E, Supplementary Fig. 2). These results indicate that dictyGrad-ALFA targeted Histone H1-mScarlet-I-ALFA for degradation in the vegetative stage of *D. discoideum.* However, even in cells expressing the standalone NbALFA (i.e., without FbxDΔWD), there was a 27% reduction in Histone H1-mScarlet-I-ALFA fluorescence (Fig. 1C, D), indicating a portion of the fluorescence reduction was not specific to dictyGrad-ALFA targeting.

**Fig. 1.**
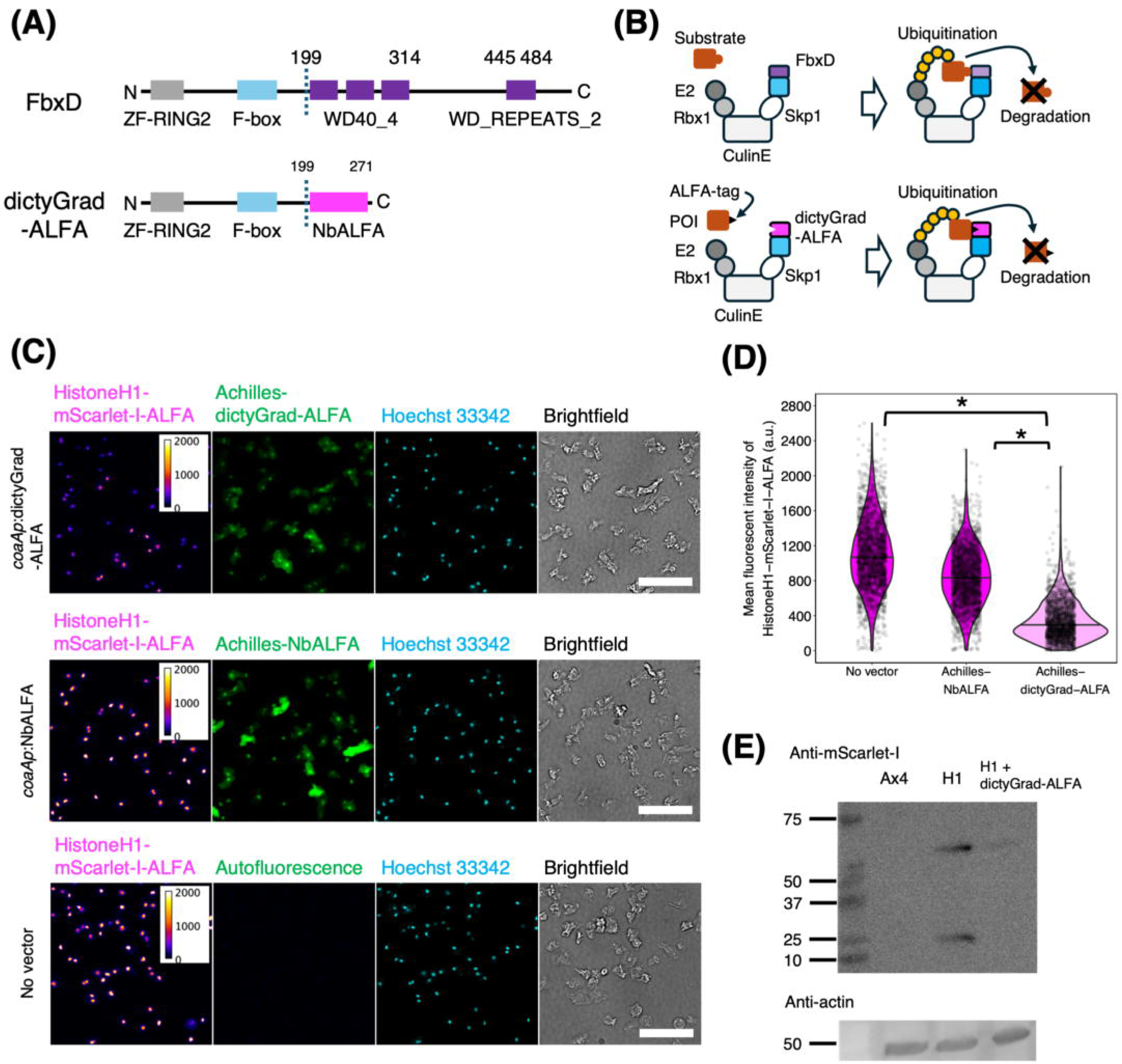
Targeted protein degradation in *D. discoideum* using dictyGrad-ALFA. (A) Domain architecture of FbxD based on InterPro annotation (upper panel) and the modified construct ‘dictyGrad-ALFA’ (lower panel). (B) Schematic overview of dictyGrad-ALFA. POI: Protein of interest. (C) Snapshots of *act5p*:Histone H1-mScarlet-I-ALFA knock-in cells carrying expression vector *coaAp*:Achilles-dictyGrad-ALFA (upper panel) or *coaAp*:Achilles-NbALFA (middle panel). No vector control (lower panel). Vegetative cells were fixed and labeled with Hoechst 33342. (Left to right) fluorescence images (heat color, Ex/Em = 561/617 nm; green, Ex/Em = 488/525 nm; cyan, Ex/Em = 405/447 nm) and brightfields. Scale bar, 50 µm. (D) Violin plots showing the mean fluorescence intensity of Histone H1-mScarlet-I-ALFA in vegetative cells. No vector (N = 2392 nuclei), Achilles-NbALFA (N = 2548 nuclei), Achilles-dictyGrad-ALFA (N = 2010 nuclei). The black horizontal line represents the median. \**P* < 10^-15^. (E) Western blots with antibodies against mScarlet-I (upper panel) and actin (lower panel). Lanes (left to right): ladder, Ax4 (parental), *act5p*:Histone H1-mScarlet-I-ALFA knock-in (H1), *act5p*:Histone H1-mScarlet-I-ALFA knock-in expressing dictyGrad-ALFA (H1 + dictyGrad-ALFA).

We then tested dictyGrad-ALFA in the developmental phase of *D. discoideum*. When dictyGrad-ALFA was expressed under the control of a prestalk-specific *ecmAO* promoter or a prespore-specific *D19* promoter, Histone-mScarlet-I-ALFA fluorescence decreased by 87% and 69% in the respective regions (the anterior 1/4 and posterior 3/4 of the slugs) than those in slugs without dictyGrad-ALFA (Fig. 2A, B). The region-specific effect was in contrast to observations in cells expressing dictyGrad-ALFA under the control of the ubiquitous *coaA* promoter, where the fluorescence intensity of Histone H1-mScarlet-I-ALFA was 98% lower throughout the slugs than that in slugs without dictyGrad-ALFA (Fig. 2C; upper panels, D). In contrast, the fluorescence intensity of Histone H1-mScarlet-I-ALFA in slugs was not affected by NbALFA expression alone (Fig. 2C; middle and lower panels, D). These results demonstrate that dictyGrad-ALFA can drive the efficient and cell type-specific degradation of constitutively expressed ALFA-tagged fluorescent proteins in the multicellular stage of *D. discoideum*.

**Fig. 2.**
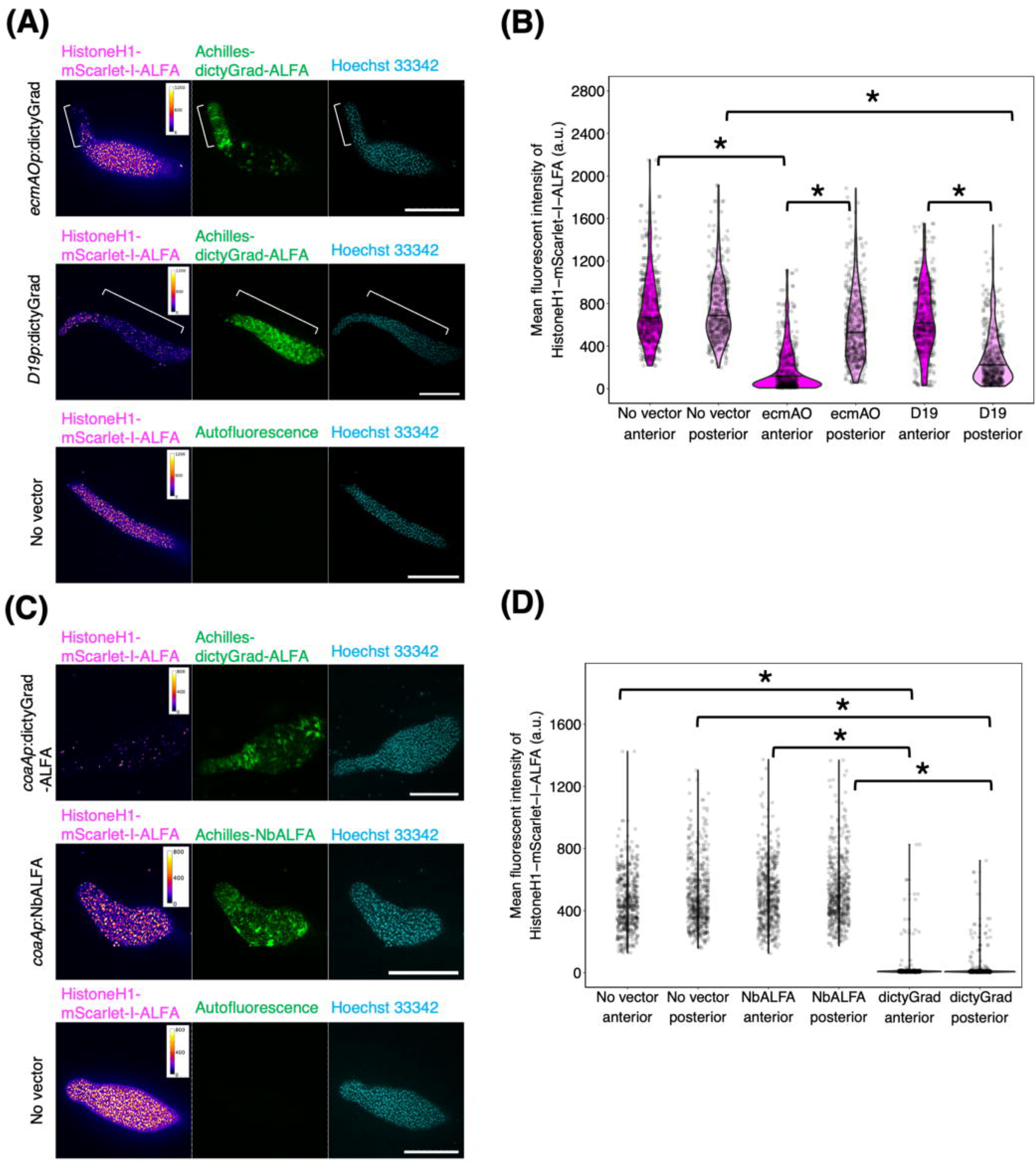
Cell type-specific protein knockdown in *D. discoideum*. (A) Knock-in strains harboring Histone H1-mScarlet-I-ALFA at the *act5* locus; Achilles-dictyGrad-ALFA expression vector driven by a prestalk-specific *ecmAO* promoter (top), prespore-specific *D19* promoter (middle), and no vector control (bottom). Slugs fixed and stained with Hoechst 33342. (Left to right) fluorescence images (hear color, Ex/Em = 561/617 nm; green, Ex/Em = 488/525 nm; cyan, Ex/Em = 405/447 nm). White brackets indicate Achilles fluorescence of the prestalk (middle) and prespore (bottom) region. The anterior-posterior axis runs from left to right. Scale bar, 100 µm. (B) Violin plots of the mean fluorescence intensity of Histone H1-mScarlet-I-ALFA. No vector control; anterior (N = 189 nuclei) and posterior (243 nuclei) parts of five slugs. ecmAO; anterior (N = 263 nuclei) and posterior (N = 448 nuclei) parts of five slugs. D19; anterior (N = 197 nuclei) and posterior (N = 466 nuclei). The black line represents the median. \**P* < 10^-15^. (C) Snapshots of fixed slugs. Knock-in strains carrying a copy of Histone H1-mScarlet-I-ALFA at the *act5* locus and the expression vector *coaAp*:Achilles-dictyGrad-ALFA (upper panel) or *coaAp*:Achilles-NbALFA (middle panel). No vector control (lower panel). Slugs were fixed and stained with Hoechst 33342. (Left to right) fluorescence images (hear color, Ex/Em = 561/617 nm; green, Ex/Em = 488/525 nm; cyan, Ex/Em = 405/447 nm). The anterior-posterior axis of the slugs runs from left to right. Scale bar, 100 µm. (D) Violin plots of the mean fluorescence intensity of Histone H1-mScarlet-I-ALFA. No vector control; anterior (N = 232 nuclei) and posterior (N = 278 nuclei) parts from four slugs. Achilles-NbALFA; anterior (N = 495 nuclei) and posterior (N = 512 nuclei) parts from four slugs. Achilles-dictyGrad-ALFA; anterior (N = 262 nuclei) and posterior (N = 344 nuclei) parts from five slugs. The black line represents the median. \**P* < 10^-15^.

### Knockdown of ACA by dictyGrad-ALFA

To further test the applicability of dictyGrad-ALFA, we targeted ACA, which is essential for the early stage of development [45] (Supplementary Fig. 3). To tag endogenous ACA, mScarlet-I-ALFA was knocked in at the 3′-end of the coding region of the *acaA* locus via CRISPR/Cas9 [4] (Fig. 3A). The knock-in by itself had no visible effect on development (Fig. 3B; ‘No dictyGrad’). ACA-mScarlet-I fluorescence was detected on the plasma membrane during the cell streaming stage, whereas the parental Ax4 cells had no detectable fluorescence (Fig. 3C; upper and lower panels). When dictyGrad-ALFA fused to the blue fluorescent protein Electra2 [46] was expressed under the constitutive *coaA* promoter, the ACA-mScarlet-I knock-in cells failed to aggregate even 24 hours after starvation (Fig. 3B; ‘+dictyGrad’). This is similar to the aggregation-less phenotype of the *acaA* knockout (Fig. 3B, Supplementary Fig 3; ‘*acaA*^-^’) [45]. After 48 hours of starvation, while most knockdown cells remained solitary, we observed a few small aggregates (Fig. 3B; ‘+dictyGrad’). The fluorescence of ACA-mScarlet-I-ALFA was almost undetectable in the majority of cells in the knockdown experiment, except for some that did not display Electra2-dictyGrad-ALFA fluorescence (Fig. 3C; middle panel). These results demonstrate that dictyGrad-ALFA mediated the efficient knockdown of ALFA-tagged ACA.

**Fig. 3.**
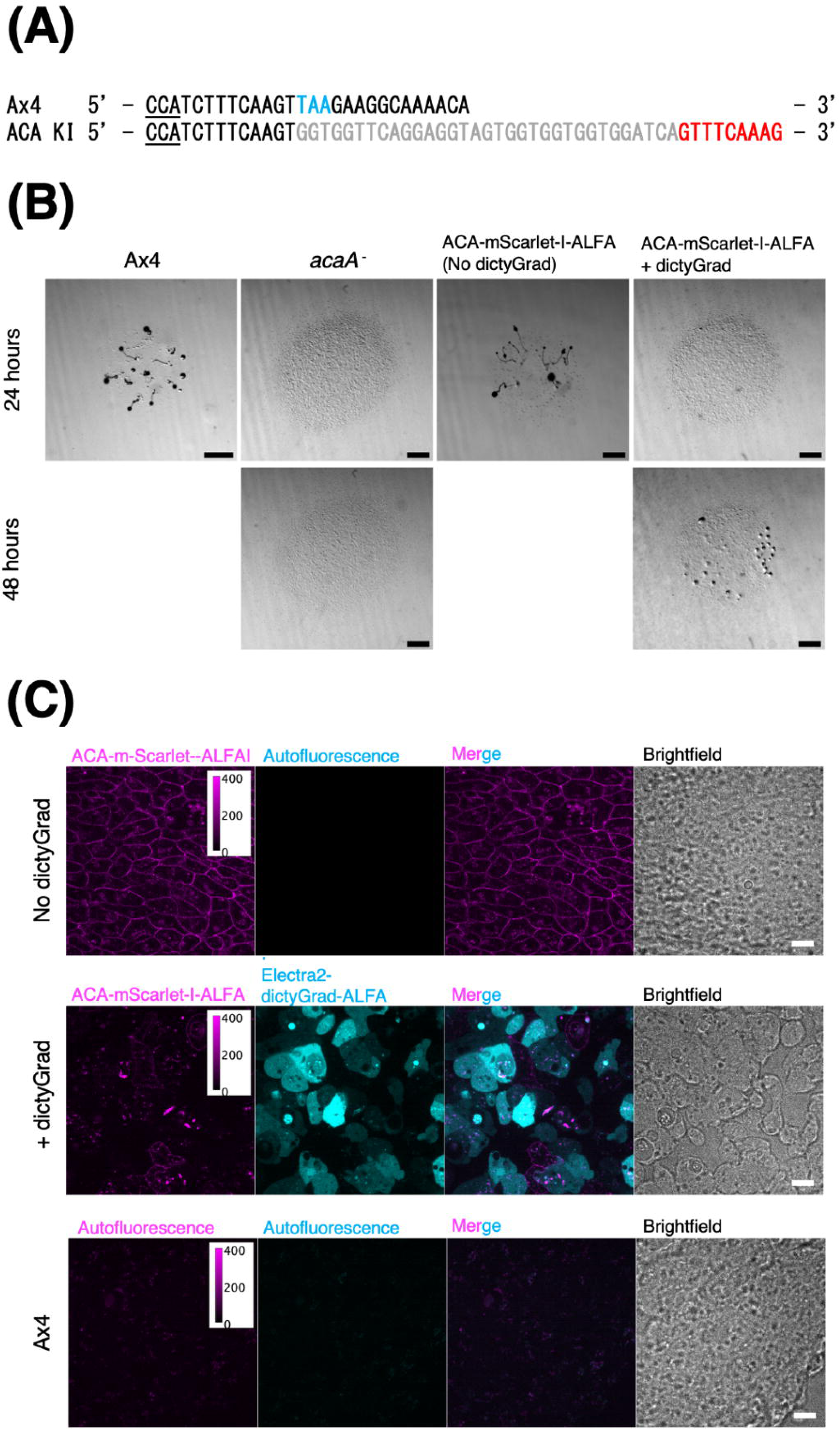
Knockdown of adenylyl cyclase ACA by dictyGrad-ALFA. (A) Diagram of the 3′ end of the *acaA* coding region in the parental Ax4 and ACA-mScarlet-I-ALFA knock-in strain. PAM sequences are underlined. Blue indicates a STOP codon. Gray and red indicate the (GGS)_2_ linker and 5′ region of mScarlet-I, respectively. (B) Transmitted light images of densely spotted cells on agar after 24 (upper panels) and 48 hours (lower panels). Strains: (Left to right) Ax4, *acaA^-^*, ACA-mScarlet-I-ALFA knock-in, and ACA-mScarlet-I-ALFA knock-in cells expressing Electra2-dictyGrad-ALFA under the *coaA* promoter. Scale bar, 1 mm. (C) Snapshots of the aggregating stream stage 7 hours after starvation. Strains: ACA-mScarlet-I-ALFA knock-in carrying no vector (top) or a vector that drives the expression of Electra2-dictyGrad-ALFA under the *coaA* promoter (middle) and Ax4 (bottom). (Left to right) fluorescence images (magenta, Ex/Em = 561/617 nm; cyan, Ex/Em = 405/447 nm), merged images, and brightfields. Scale bar, 10 µm.

We next examined whether the cell type-specific degradation of ACA affects development in the multicellular stage. The parental Ax4 and ACA-mScarlet-I-ALFA knock-in strain formed fruiting bodies containing stalk cells and spores with cellulose-coated cell walls 48 hours after starvation (Fig. 4A and B). Strains that did not carry dictyGrad-ALFA formed fruiting bodies by 24 hours (Fig. 4A; ‘Ax4’ and ACA-mScarlet-I-ALFA knock-in ‘No dictyGrad’), and ACA knock-in cells expressing dictyGrad-ALFA under the control of the prestalk or prespore promoter showed marked delays in development and remained at the slug stage (Fig. 4A; 24 hours). By 48 hours, these strains completed culmination (Fig. 4A). The morphologies of stalk cells and spores of these knockdown strains were indistinguishable from those of the controls (Fig. 4B). Accordingly, dictyGrad-ALFA expression in the prestalk or prespore cells decreased the fluorescence signals of ACA-mScarlet-I in their respective slug regions (Fig. 5A, B). These patterns contrast with the ACA-mScarlet-I fluorescence patterns observed throughout the slugs of the ACA-mScarlet-I-ALFA knock-in strain (Fig. 5A, B). Note that, similar to aggregation-stage cells, ACA-mScarlet-I appeared as perforated clusters in the plasma membrane of the slug stage cells (Fig. 5B). The small aggregates in the cytosol were autofluorescent (Fig. 5B). In the fruiting bodies, ACA-mScarlet-I fluorescence at the plasma membrane was visible mostly in stalk cells (Fig. 5C, D). It was detected at the plasma membrane and in intracellular particles in control (No dictyGrad-ALFA) strains and in cells expressing dictyGrad-ALFA under the control of the *D19* promoter (Fig. 5C, D). In contrast, when dictyGrad-ALFA was expressed under the control of the *ecmAO* promoter, ACA-mScarlet-I fluorescence was markedly reduced in the plasma membrane (Fig. 5C, D). These results indicate that while cell-type-specific ACA knockdown delayed development substantially, it had no apparent effect on fruiting body formation.

**Fig. 4.**
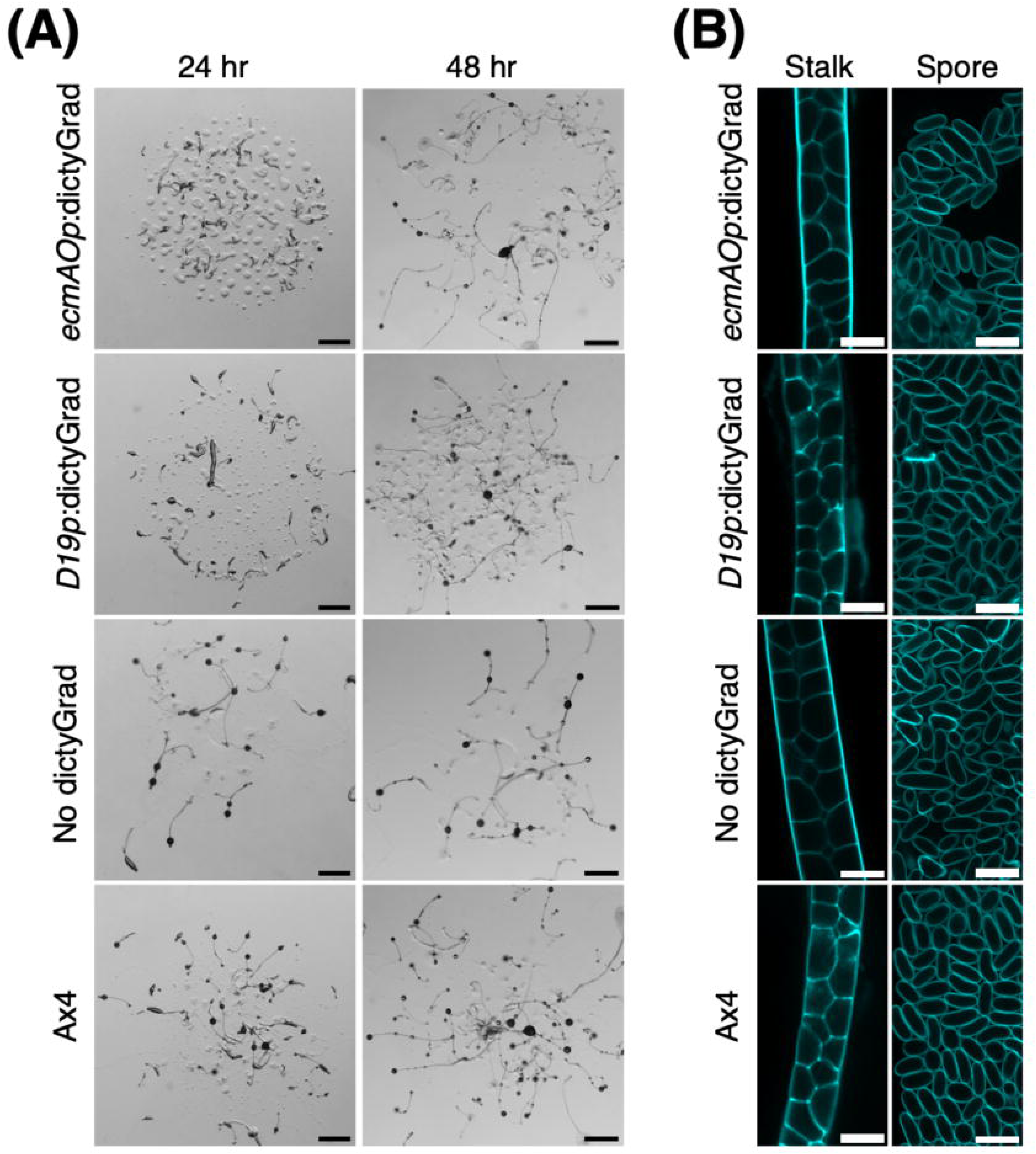
Cell type-specific knockdown of ACA delays development. (A) Snapshots of developing ACA-mScarlet-I-ALFA knock-in strains on agar plates 24 hours after starvation. Scale bar, 1 mm. (B) Fluorescence images (Ex/Em = 405/447 nm: left) of stalk tubes (left panel) and spores (right panel) 48 hours after starvation. Cellulose in cell walls was labelled with calcofluor. Scale bar, 10 µm. Top to bottom; Achilles-dictyGrad-ALFA expression vector driven by the prestalk-specific *ecmAO* promoter, prespore-specific *D19* promoter, no dictyGrad vector, and Ax4.

**Fig. 5.**
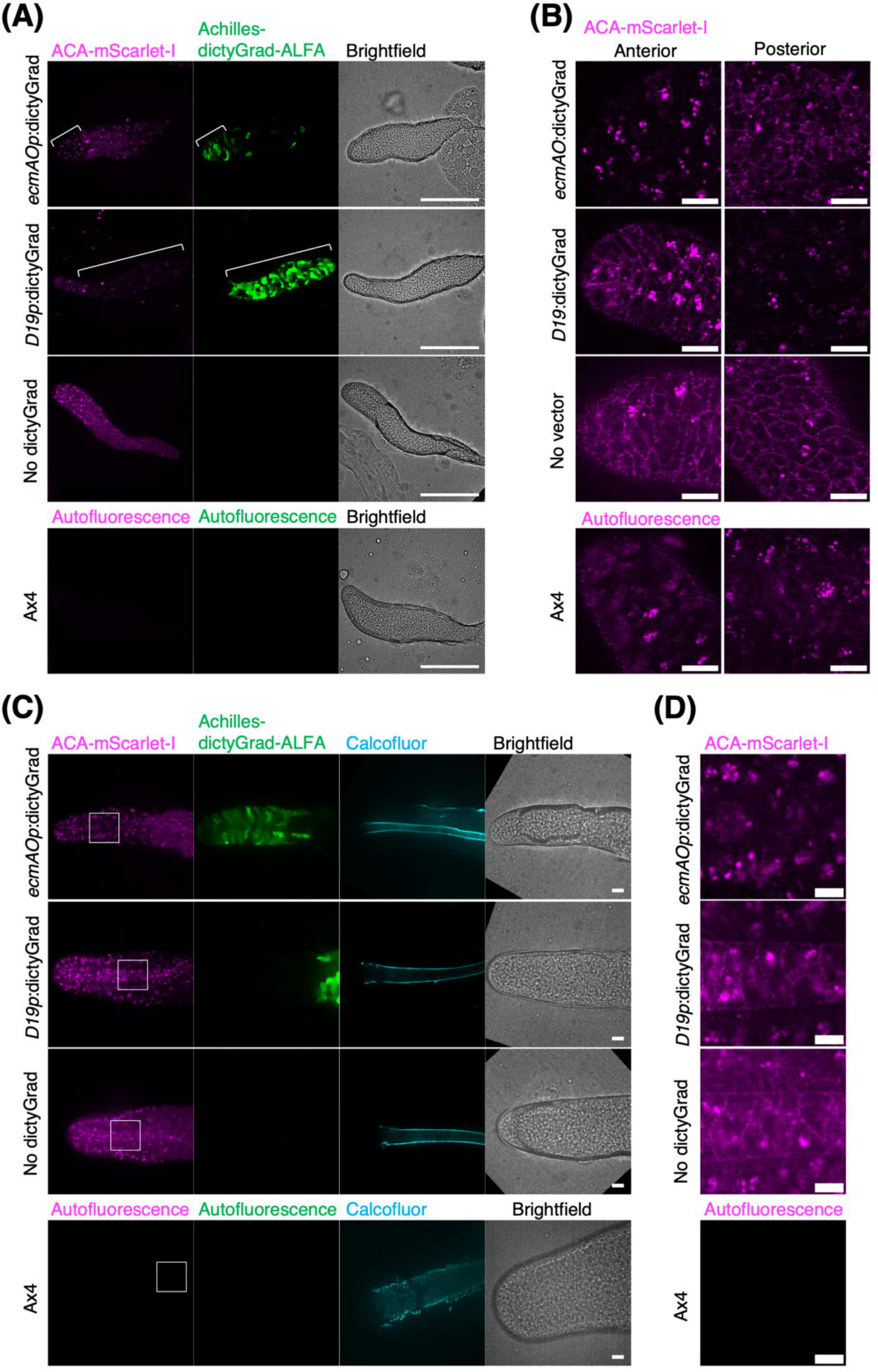
Cell type-specific reduction of ACA via dictyGrad-ALFA. (A) Snapshots of the slug stage 20 hours after starvation. (Top to bottom) Achilles-dictyGrad-ALFA expression vector driven by the prestalk-specific *ecmAO* promoter, prespore-specific *D19* promoter, no dictyGrad vector, and Ax4. (Left to right) fluorescence images (magenta, Ex/Em = 561/617 nm; green, Ex/Em = 488/525 nm), and brightfields. Scale bar, 100 µm. (B) Higher magnification ACA-mScarlet-I-ALFA images of the anterior (left panel) and posterior region (right panel) of slugs shown in (A). Scale bar, 10 µm. (C) Snapshots of culminants observed 24 hours after starvation. Cellulose in stalk tubes was labelled with calcofluor. (Top to bottom) Achilles-dictyGrad-ALFA expression vector driven by the prestalk-specific *ecmAO* promoter, prespore-specific *D19* promoter, no dictyGrad vector, and Ax4. (Left to right) fluorescence images (magenta, Ex/Em = 561/617 nm; green, Ex/Em = 488/525 nm; cyan, Ex/Em = 405/447 nm), and brightfields. Scale bar, 10 µm. (D) Higher magnification ACA-mScarlet-I-ALFA images of stalk tubes shown by white boxes in (C). Scale bar, 10 µm.

### Application of the AID2 system to *D. discoideum* cells

Next, we incorporated the AID2 (auxin-inducible degron2) system for conditional protein degradation. AID2 employs a mAID tag for the targeted degradation of a POI, TIR from *Oryza sativa* with the F74G mutation (OsTIR(F74G)) and a synthetic auxin analog 5-Phenyl-1H-indole-3-acetic acid (5-Ph-IAA), which mediates dimerization between the POI and OsTIR(F74G) (Fig. 6A) [24]. The presence of 10 µM 5-Ph-IAA in the liquid medium had a negligible effect on the growth rate of *D. discoideum* (10.02 ± 0.44 hours with DMSO and 9.95 ± 0.91 hours with 10 µM 5-Ph-IAA, three experiments: Supplementary Fig. 4A). Drawing on an example in mammalian cell lines where GFP-tagged proteins were targeted by an anti-GFP nanobody fused to mAID [47], we employed a Histone H1-mScarlet-I-ALFA knock-in strain that co-expressed mAID-NbALFA and a fusion protein of *D. discoideum* Skp1A and OsTIR(F74G)-Achilles (Fig. 6A). The Skp1 fusion in this construct was based on a scheme reported for budding yeast [48]. When treated with 5-Ph-IAA during growth for 24 hours, there was a 60% decrease in Histone H1-mScarlet-I fluorescence compared with that in the mock DMSO control (Fig. 6B, C). Western blotting against mScarlet-I revealed 78% and 58% reductions in Histone H1-mScarlet-I levels with or without IAA (Fig. 6D, Supplementary Fig. 6).　While this equates to an IAA-induced reduction of approximately 50% in the mScarlet-I fluorescence intensity, there was significant background degradation. A similar but slightly weaker mScarlet-I fluorescence change was observed for OsTIR(F74G)-Achilles without the Skp1A fusion (Supplementary Fig. 5).

**Fig. 6.**
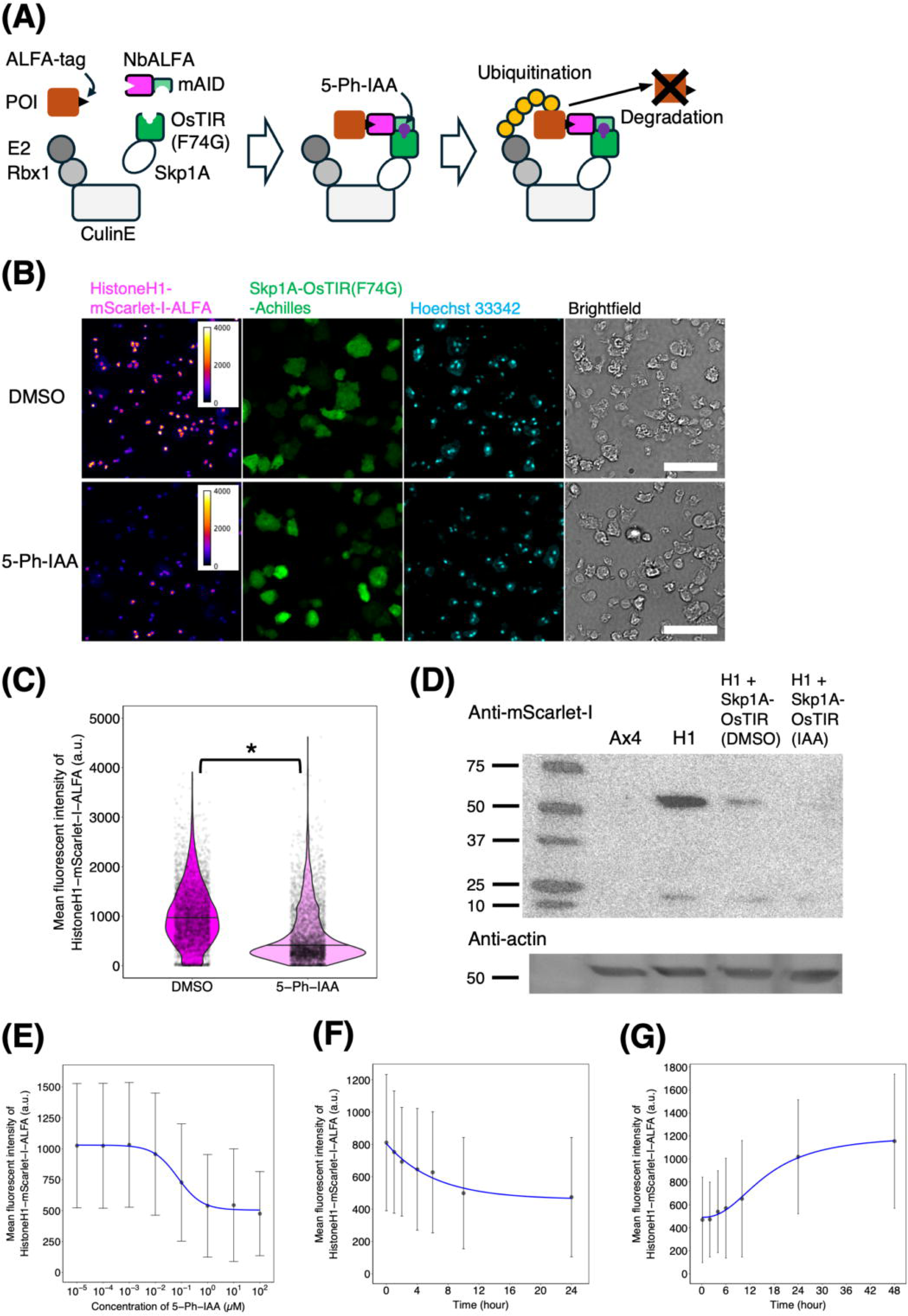
IAA-inducible degron in *D. discoideum*. (A) Schematic overview of the ALFA tag-based auxin-inducible degron. POI: Protein of interest. (B) Snapshots of *act5p*:Histone H1-mScarlet-I-ALFA knock-in cells carrying the expression plasmids *coaAp*:Skp1A-OsTIR(F74G)-Achilles and *act15p*:mAID-NbALFA. Vegetative cells were fixed and stained with Hoechst 33342. (Left to right) fluorescence images (hear color, Ex/Em = 561/617 nm; green, Ex/Em = 488/525 nm; cyan, Ex/Em = 405/447 nm) and brightfields. DMSO (upper panel) and 10 µM 5-Ph-IAA (lower panel) treatment. Scale bar, 50 µm. (C) Violin plots of the mean fluorescence intensity of Histone H1-mScarlet-I-ALFA for DMSO (N = 3771 nuclei) and 5-Ph-IAA (N = 3961 nuclei). The black line represents the median. \**P* < 10^-15^. (D) Western blots with antibodies to mScarlet-I (upper) and actin (lower). Lanes (left to right): ladder, Ax4, Histone H1-mScarlet-I-ALFA *act5* locus knock-in (H1), a Histone H1-mScarlet-I-ALFA *act5* locus knock-in carrying the expression vector *coaAp*:Skp1A-OsTIR(F74G)-Achilles treated for 24 hours with either DMSO (H1 + Skp1A-OsTIR (DMSO)) or 10 µM 5-Ph-IAA (H1 + Skp1A-OsTIR (IAA)). (E) 5-Ph-IAA dose-response of the fluorescence intensity of Histone H1-mScarlet-I-ALFA after 24 hours (H1 + Skp1A-OsTIR cells). Mean ± SD; 0.01 nM (N = 6315 nuclei), 0.1 nM (N = 6190 nuclei), 1 nM (N = 5534 nuclei), 10 nM (N =5556 nuclei), 100 nM (N = 5524 nuclei), 1 µM (N = 4794 nuclei), 10 µM (N = 4889 nuclei), 100 µM (N = 9724 cells). (F) Decrease in the fluorescence intensity of Histone H1-mScarlet-I-ALFA following the addition of 10 µM 5-Ph-IAA. Mean ± SD; 0 hours (N = 1438 nuclei), 1 hour (N = 3964 nuclei), 2 hours (N = 3676 nuclei), 4 hours (N = 4294 nuclei), 6 hours (N = 3134 nuclei), 10 hours (N = 5465 nuclei), 24 hours (N = 5465 nuclei). (G) Recovery of the fluorescence intensity of Histone H1-mScarlet-I-ALFA after the removal of 10 µM 5-Ph-IAA. Cells were treated with 5-Ph-IAA for 24 hours before medium exchange. Mean ± SD; 0 hours (N = 4567 nuclei), 2 hours (N = 6467 nuclei), 4 hours (N = 6354 nuclei), 6 hours (N = 8053 nuclei), 10 hours (N = 2047 nuclei), 24 hours (3547 nuclei), 48 hours (N = 3724 nuclei).

In analyses of dose-dependency and kinetics of ALFA-tagged Histone H1-mScarlet-I fluorescence, the DC_50_ value, representing the 5-Ph-IAA concentration required to reduce the average fluorescence intensity by 50% after 24 hours, was 72 nM (Fig. 6E). This was approximately 10–100 times higher than those observed in HCT116 cells [24] and *Caenorhabditis elegans [25]*. The time until 50% reduction (i.e., half-life) under 10 µM 5-Ph-IAA treatment was 4.35 hours (Fig. 6F), which was 4 to 15 times longer than those reported for HCT116 cells [24] and *C. elegans* larvae [25]. When we examined the recovery of Histone H1-mScarlet-I fluorescence after removing 5-Ph-IAA, the half-time was approximately 15.9 hours (Fig. 6G), which was 8 times longer than that reported for HCT116 cells *[24]*. These results indicated a low efficiency of 5-Ph-IAA-induced targeted protein degradation in the growth stage of *D. discoideum*. As expected, when the ACA-mScarlet-I knock-in cells expressing Skp1A-OsTIR-Achilles under the *coaA* promoter were starved and submerged in buffer with 10 µM 5-Ph-IAA, ACA-mScarlet-I-ALFA fluorescence was still visible at the plasma membrane at a comparable level to that in the mock control (Supplementary Fig. 7). Accordingly, cell aggregation was observed (Supplementary Fig. 7).

We further examined AID2-mediated protein degradation in the multicellular stage of *D. discoideum*. The inclusion of 10 µM 5-Ph-IAA in the filter pad had no apparent effect on wild-type development (Supplementary Fig. 4B). In the slugs developed on a filter pad containing IAA, fluorescence intensity of Histone H1-mScarlet-I expressed under the constitutive *act5* promoter was reduced by 23% in cells at the posterior region and by 65% at the anterior region (Fig. 7A, B, C). While the fluorescence intensity of Skp1A-OsTIR(F74G)-Achilles was relatively high in the anterior region (Fig. 7B, C), we did not find a strong correlation with Histone H1-mScarlet-I fluorescence (Fig. 7D, E). These results suggest that while the AID system can support inducible protein degradation in prestalk cells, prespore cells are more resistant to the perturbation.

**Fig. 7.**
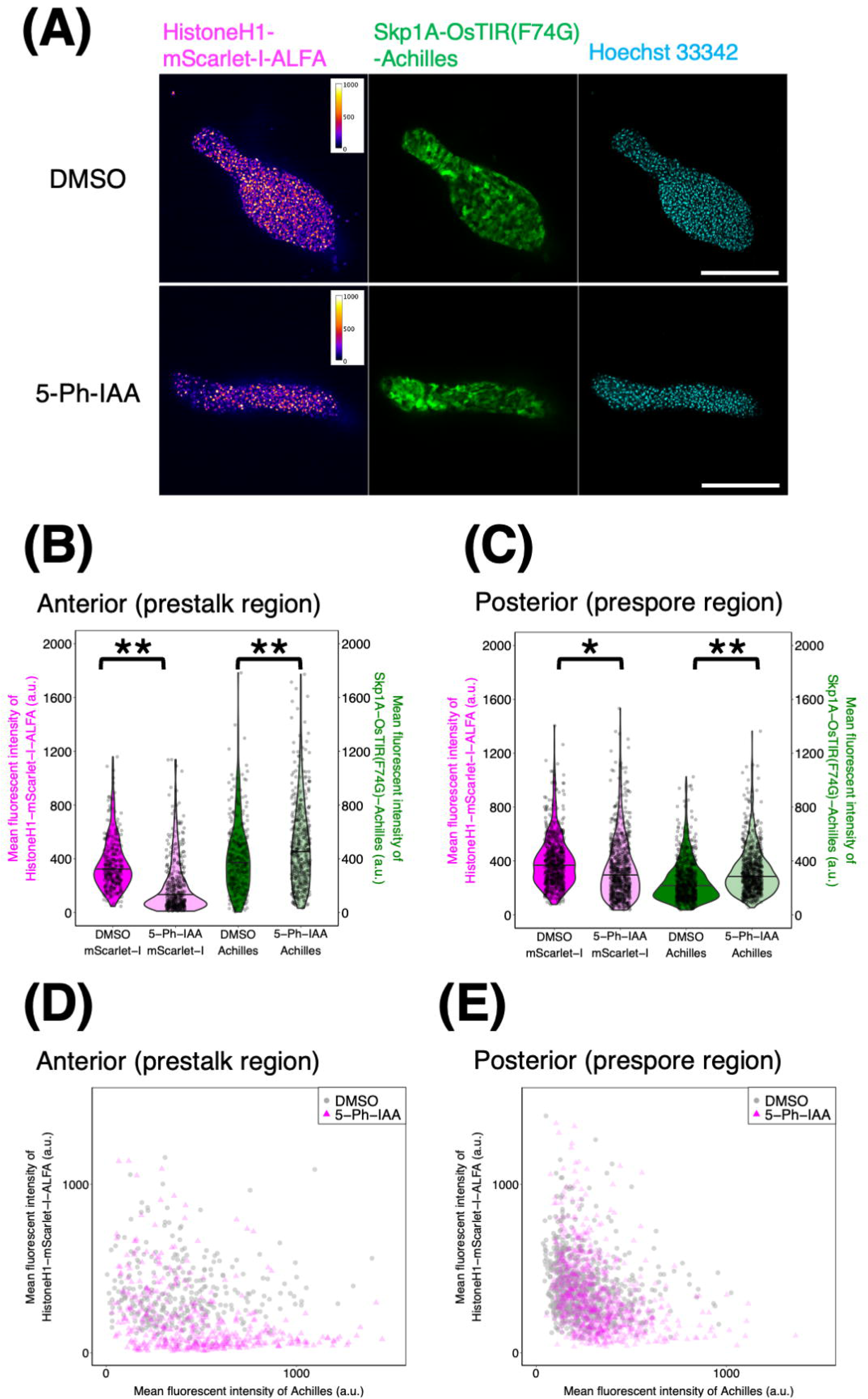
IAA-induced protein degradation in the slug stage. (A) Snapshots of a Histone H1-mScarlet-I-ALFA *act5* locus knock-in carrying the expression vector *coaAp*:Skp1A-OsTIR(F74G)-Achilles. The anterior-posterior axis of the slugs runs from left to right. (Left to right) Fluorescence images (magenta, Ex/Em = 561/617 nm; green, Ex/Em = 488/525; cyan, Ex/Em = 405/447 nm). Cells were developed for 20 hours with 10 µM 5-Ph-IAA (lower panel) or with a mock control (DMSO; upper panel), fixed, and stained with Hoechst 33342. Scale bar, 100 µm. (B, C) Violin plots showing the mean fluorescence intensity of nuclear Histone H1-mScarlet-I-ALFA (magenta) and Skp1A-OsTIR(F74G)-Achilles (green) in anterior (B) and posterior (C) part of slugs. For both (B) and (C): DMSO–treated; anterior (N = 305 nuclei) and posterior (N = 712 nuclei) regions from five slugs. 5-Ph-IAA–treated; anterior (N = 430 nuclei) and posterior (N = 770 nuclei) regions from five slugs. The black line represents the median. \*\**P* < 10^-15^. \**P* < 10^-12^. (D, E) The mean fluorescence intensity of Histone H1-mScarlet-I-ALFA plotted against that of Achilles fused to Skp1A-OsTIR(F74G) in anterior (D) and posterior (E) part of slugs (Anterior–DMSO, N = 305 nuclei. Anterior–IAA, N = 430 nuclei. Posterior–DMSO, N = 712 nuclei. Posterior–IAA, N = 770 nuclei).

## Discussion

In this study, we established a protein knockdown system in *D. discoideum* based on a nanobody-based degron. The expression of dictyGrad-ALFA led to a marked decrease in the fluorescence signal of Histone-mScarlet-I-ALFA in vegetative cells. Cell type-specific Histone-mScarlet-I-ALFA knockdown was achieved by driving dictyGrad-ALFA expression under a prestalk- or prespore-specific promoter. Furthermore, we demonstrated the knockdown of ACA, which is essential for cell aggregation in the early stage of *D. discoideum* development. The fluorescence of ACA-mScarlet-I-ALFA was not detectable under the constitutive expression of dictyGrad-ALFA, leading to the expected aggregation-null phenotype.

The use of ALFA-tag to target a POI to a nanobody-F-box complex leaves GFP and other conventional fluorescent markers in the green spectra available for downstream analyses. This is especially useful for analyses requiring targeted knock-down combined with multi-color imaging [30]. In addition, the ALFA-tag, due to its small size, is expected to minimize interference with the function of endogenous proteins, thereby facilitating knock-down studies of sensitive loci where the addition of fluorescent tags alone has adverse effects [49]. Alternatively, a GFP tag-based approach may be useful for studies using GFP-knock-in strains that are already available. Although not tested, replacing NbALFA with an anti-GFP nanobody should, in principle, support such knockdown studies in *D. discoideum*. Other short peptide tags, such as BC2-tag, EPEA-tag, MoonTag, and PepTag, and their nanobodies [50–54] should support the simultaneous or context-dependent selective degradation of multiple target proteins in the future. Using various promoters as well as a Dox-on inducible system [55], dictyGrad-ALFA should enable cell type- and stage-specific conditional knockdown studies, which have not been possible in *D. discoideum* to date.

Using the ACA-mScarlet-I-ALFA knock-in strain, ACA localized at the plasma membrane in both streaming- and slug-stage cells (Fig. 3, 5), consistent with earlier observations of cell streams of a ACA-GFP knock-in strain [56] and immunostaining results for isolated slug cells [57] but dissimilar to the strong posterior accumulation of overexpressed YFP-tagged ACA [58]. While an earlier immunostaining study suggested that ACA is expressed specifically in prestalk cells [57], we observed ACA-mScarlet-I-ALFA fluorescence in both prestalk and prespore cells (Fig. 5). This is not surprising given that both prestalk and prespore cells are capable of a ‘cAMP relay response’ (i.e., G-protein-coupled receptor-mediated synthesis of cAMP in response to external cAMP stimuli) [59, 60]. Since *ecmAO* and *D19* promoter activity occurs during the streaming stage and increases thereafter [30], the delayed development suggests that ACA likely plays a role in directing cell movements and/or cell differentiation during the mound stage. Plasma membrane-bound ACA was mostly visible in stalk cells during the culmination phase (Fig. 5); however, its loss had no obvious effect on culmination (Fig. 4, 5), suggesting that ACA is dispensable for fruiting body formation. While this is counter to the notion that cyclic-di-GMP-induced cAMP synthesis via ACA is necessary for stalk gene induction through PKA activation [61], it is in line with the observation that *acaA* knockout mutants pretreated with cAMP pulses show agglutination and culmination [60, 62].

We should note that the degradation of ALFA-tagged Histone H1-mScarlet-I was incomplete (Fig. 1). This may be due to its excessive expression driven by the strong *act5* promoter, resulting in a protein level exceeding the degradation capacity. While we expect that most native proteins are not expressed at this high level, a more efficient system capable of degrading highly expressed proteins may be achieved using an alternative F-box domain, such as those from FbxA [63] and other F-box proteins in *D. discoideum*. The small aggregates of ACA-mScarlet-I-ALFA knock-in cells expressing dictyGrad-ALFA observed after 48 hours (Fig. 3) may be due to heterogeneity in nanobody expression levels. Although a clonal strain was analyzed, because the nanobody was expressed from an extrachromosomal plasmid, variation in the plasmid copy number may have contributed to this heterogeneity. Even a small population of cells with ineffective knockdown may mask the developmental phenotype due to a cell non-autonomous effect. To minimize cells with reduced nanobody expression, it may be effective to employ random integration *[64, 65]* to generate a strain with multiple copies of dictyGrad-ALFA in the genome.

For conditional knockdown, we combined our ALFA tag-based approach in *D. discoideum* with the AID2 system. We tested whether an ALFA-tagged protein can be targeted for degradation in *D. discoideum* cells by expressing *O. sativa* TIR and an anti-ALFA nanobody fused to mAID [23, 24]. We observed the 5-Ph-IAA-induced degradation of Histone H1-mScarlet-I-ALFA using a strain expressing OsTIR(F74G) with or without Skp1A fusion (Fig. 6, Supplementary Fig. 5). Several issues must be addressed, however, for the practical use of the AID system in *D. discoideum*. Although the IAA modification and mutations in the TIR binding pocket in the AID2 system were developed for low basal degradation [24], we detected a high basal degradation rate, apparent low potency of 5-Ph-IAA, and slow degradation and recovery rates. These factors likely contributed to the lower efficiency than other model systems, such as HCT116 cells and *C. elegans* [24, 25]. Although dictyostelids produce various natural compounds [66, 67] and IAA degradation is known in plant-associated bacteria [68], there is no evidence that *D. discoideum* can metabolize indole. Active export by shedding extracellular vesicles [69] and ABC transporters [70] have been reported in *D. discoideum* and thus may have contributed to the requirement for a high dosage of 5-Ph-IAA. We should note that OsTIR(F74G) expression gave rise to high background protein degradation (i.e., without IAA) (Fig. 6D). This may be attributed to cross-talk between OsTIR/mAID and endogenous factors. Since prestalk cells were more susceptible to IAA induction, it is possible that potential cross-talk and/or IAA exclusion is relatively limited in this cell type. The issue can potentially be circumvented by the recently reported analog 5-Ad-IAA, which binds to another mutant form, OsTIR(F74A), with higher specificity and targeting efficiency in vertebrate cells [71]. Alternatively, other tools that mediate dimerization either chemically [72] or optically [73] may mitigate the issue.

## Conclusions

We established a nanobody-based degron system for knockdown studies in *D. discoideum*. Expression of the F-box domain fused to an anti-ALFA tag-nanobody (dictyGrad-ALFA) directs efficient targeted protein degradation and should facilitate stage- and cell type-specific loss-of-function analyses. Although less efficient, the auxin-inducible degron (AID2) system may be useful in analyzing genes essential for cell survival. These systems are expected to serve as powerful tools for analyzing the roles of essential genes in development and survival in *D. discoideum*.

## Declarations

### Ethics approval and consent to participate

Not applicable.

### Consent for publication

Not applicable.

### Availability of data and materials

The datasets used and/or analyzed during the current study are available from the corresponding author on reasonable request.

### Competing interests

The authors declare no competing interests.

### Funding

This work was supported by JST CREST JPMJCR1923, JSPS KAKENHI JP19H05801, JP23H00384, JP25H01360, JP25H01771, JP25K22490 grants and HFSP Research Grant RGP0051/2021 to SS. JP20J00751, JP21K15081, and JP23H04304 grants to HH; JP22K15119 grant to SK.

## Acknowledgments

We thank Douwe Veltman for pDM304 (National BioResource Project Nenkin plasmid; G90008), Tetsuya Muramoto for pTM1285 and pTM1224 (National BioResource Project Nenkin plasmid; G90246 and G90427, respectively), Taihei Fujimori for technical support. The plasmids constructed in this study will be available from the NBRP-Nenkin stock center (https://nenkin.nbrp.jp/).

## Author information

### Contributions

H.H. and S.S. conceived the study. H.H. designed the schemes, acquired data and performed analysis. H.H., S.F., T.A., T.S. constructed plasmids. S. K. generated *acaA* null strain. N.S performed western blot analysis. H.H. and S.S. wrote the manuscript.

## Notes

### Competing Interest Statement

The authors have declared no competing interest.

