## Supplementary Information for "A nanobody-based degron system for targeted protein knockdown in *Dictyostelium discoideum*"

### **Supplementary materials**

#### **Sequences synthesized in this study**

>ALFA-tag

CCATCAAGATTAGAAGAAGAATTACGTCGTCGTTTAACAGAACCA

> NbALFA

ATGGAAGTACAATTACAAGAATCAGGAGGAGGTTTAGTTCAACCAGGTGGTTCATTAAGATTAAGTTGTACAGCTAGTGGTGTAACAATTAGTGCATTAAATGCTATGGCCATGGGTTGGTATAGACAAGCTCCAGGTGAAAGAAGAGTAATGGTTGCAGCAGTTTCTGAAAGAGGTAATGCAATGTATCGTGAATCTGTTCAAGGTAGATTTACTGTTACAAGAGATTTTACCAATAAAATGGTCTCATTGCAAATGGATAATTTGAAACCAGAAGATACTGCAGTTTATTATTGTCATGTATTAGAGGATCGTGTAGATTCATTTCATGATTATTGGGGTCAAGGTACACAAGTTACTGTTTCATCATAA

>mAID

AAGGAAAAGTCTGCATGTCCAAAAGATCCAGCTAAACCACCAGCAAAAGCACAAGTAGTTGGTTGGCCACCTGTTCGTAGTTACAGAAAGAATGTTATGGTTTCATGTCAAAAGTCATCAGGTGGTCCAGAAGCCGCTGCATTTGTTAAGGTTTCAATGGATGGTGCCCCATACTTAAGAAAGATTGATTTGAGAATGTATAAA

> OsTIR

ATGACCTACTTCCCAGAAGAAGTTGTTGAGCATATCTTCTCATTCTTGCCTGCTCAAAGAGATCGTAATACTGTATCTTTGGTTTGCAAAGTTTGGTATGAAATTGAGCGTTTGTCAAGAAGAGGAGTTTTCGTTGGAAACTGCTACGCCGTTAGAGCTGGTCGTGTTGCCGCAAGATTTCCTAATGTACGTGCTTTGACAGTTAAGGGTAAGCCTCACTTCGCCGATTTTAACTTGGTTCCTCCAGATTGGGGAGGTTACGCCGGACCTTGGATAGAGGCAGCTGCAAGAGGTTGTCACGGTTTGGAAGAACTTAGAATGAAGCGTATGGTAGTTTCTGATGAAAGTCTTGAACTTTTAGCACGTTCATTCCCTAGATTTCGTGCATTAGTACTTATTTCATGTGAAGGATTTTCAACTGATGGTTTGGCCGCTGTAGCTTCACACTGTAAATTGCTTAGAGAATTAGATTTGCAAGAAAACGAGGTTGAAGATCGTGGTCCACGTTGGTTGTCTTGCTTTCCAGATTCTTGCACCTCATTAGTAAGTTTGAACTTTGCTTGCATTAAGGGAGAAGTTAACGCAGGAAGTCTTGAGCGTTTAGTTTCTCGTAGTCCTAACTTACGTTCACTTAGATTGAACCGTAGTGTTTCAGTTGATACTTTGGCCAAAATCCTTTTGCGTACACCTAACCTTGAAGATTTAGGAACTGGAAACTTGACCGATGATTTTCAAACAGAAAGTTACTTCAAGTTGACTTCTGCTTTGGAGAAGTGCAAAATGCTTAGATCATTATCAGGATTCTGGGATGCTTCTCCTGTATGCTTGAGTTTCATTTACCCTTTGTGTGCTCAATTGACCGGATTGAATCTTTCATACGCCCCTACATTGGATGCATCTGATTTAACTAAAATGATCTCTCGTTGCGTAAAATTGCAACGTTTGTGGGTATTGGATTGCATCTCAGATAAGGGTTTGCAAGTTGTAGCAAGTAGTTGCAAGGATTTACAAGAGTTGCGTGTATTCCCATCTGATTTCTACGTAGCAGGATACTCTGCAGTTACTGAGGAAGGTTTGGTTGCTGTATCTTTAGGATGTCCTAAGTTAAATAGTTTGTTATACTTTTGCCACCAAATGACTAATGCAGCCTTGGTAACCGTTGCTAAGAACTGCCCTAACTTTACAAGATTCAGATTATGCATTTTAGAACCTGGTAAACCTGATGTTGTAACTTCACAACCACTTGATGAAGGTTTCGGAGCCATAGTACGTGAATGCAAGGGTTTACAAAGACTTAGTATCAGTGGTTTGTTAACCGATAAGGTATTCATGTACATTGGAAAGTACGCTAAGCAACTTGAGATGTTATCTATAGCTTTCGCAGGAGATTCTGATAAGGGAATGATGCATGTTATGAATGGATGTAAGAACTTAAGAAAGTTAGAGATTCGTGATTCACCATTCGGAGATGCCGCTTTGTTAGGAAATTTCGCAAGATACGAAACAATGCGTAGTTTATGGATGTCTTCTTGTAACGTAACACTTAAGGGATGCCAAGTATTAGCTTCTAAGATGCCAATGTTAAATGTTGAGGTTATAAATGAGCGTGATGGATCTAATGAGATGGAAGAGAATCATGGAGATTTACCTAAGGTTGAAAAGTTGTATGTTTATCGTACTACCGCCGGTGCACGTGATGATGCTCCAAACTTCGTAAAGATTTTATAA

#### **Supplementary Table 1. List of plasmids used in this study**

| Plasmid index | Plasmid name | Related figure | Reference |
| --- | --- | --- | --- |
| – | pDM304 | – | [1] |
| #HH498 | pDM304_*act15p*:HistoneH1-mTagBFP2 | Supplementary Fig. 1 | [2] |
| #HH570 | pDM304_MCS-(GGS)_2_-mTagBFP2-ALFA | Supplementary Fig. 1 | This study |
| #HH573 | pDM304_*act15p*:HistoneH1-mTagBFP2-ALFA | Supplementary Fig. 1 | This study |
| #HH242 | pDM358_*act15p*:mScarlet-I-(GGS)_2_-MCS | – | [2] |
| #HH574 | pDM358_*act15p*:mScarlet-I-NbALFA | Supplementary Fig. 1 | This study |
| #HH608 | pDM1501_HistoneH1-mScarlet-I-ALFA | Fig. 1, 2, 6, 7 | This study |
| #HH162 | pDM326_*act15p*:Achilles-(GGS)_2_-MCS | – | [2] |
| #HH623 | pDM326_*act15p*:Achilles-NbALFA | Fig. 1 | This study |
| #HH109 | pDM304_*coaAp*:PH_Akt_-mNeonGreen | – | [2] |
| #HH707 | pDM304_*coaAp*:Achilles-NbALFA | Fig. 1, 2 | This study |
| #HH708 | pDM304_*coaAp*:Achilles-FbxDΔWD-NbALFA | Fig. 1, 2 | This study |
| #HH657 | pDM304_*ecmAOp*:Achilles-FbxDΔWD-NbALFA | Fig. 2, 4, 5 | This study |
| #HH659 | pDM304_*D19p*:Achilles-FbxDΔWD-NbALFA | Fig. 2, 4, 5 | This study |
| #HH676 | pDM304_*act15p*:mAID-NbALFA | Fig. 6, 7 | This study |
| #HH697 | pDM304_*coaAp*:OsTIR(F74G)-Achilles | Supplementary Fig. 5 | This study |
| #HH699 | pDM304_*coaAp*:OsTIR(F74G)-Achilles_*act15p*:mAID-NbALFA | Supplementary Fig. 5 | This study |
| #HH253 | pDM358_*act15p*: MCS-(GGS)_2_-mScarlet-I | – | [2] |
| #HH350 | pDM358_*fpaAp*:Skp1A-mScarlet-I | Fig. 6, 7 | This study |
| #HH698 | pDM304_*coaAp*:Skp1A-(GGS)_2_-OsTIR(F74G)-Achilles | Fig. 6, 7, Supplementary Fig. 7 | This study |
| #HH700 | pDM304_*coaAp*:Skp1A-(GGS)_2_-OsTIR(F74G)-Achilles_*act15p*:mAID-NbALFA | Fig. 6, 7, Supplementary Fig. 7 | This study |
| – | pTM1285 | – | [3] |
| #HH681 | pTM1285_*acaA*(4514-4534) | Fig. 3 | This study |
| – | pTM1224 | Supplementary Fig. 3 | [3] |
| #HH722 | pDM358_*act15p*:ACA-mScarlet-I | Supplementary Fig. 3 | This study |
| #HH686 | pDM304_*coaAp*:Electra2-NbALFA | Fig. 3 | This study |
| #HH687 | pDM304_*coaAp*:Electra2-FbxDΔWD-NbALFA | Fig. 3 | This study |

* (GGS)_2_: Gly-Gly-Sr-Gly-Gly-Ser linker

#### **Supplementary Table 2. List of primers used in this study**

| Primer name | Sequence (5′–3′) | Related plasmid |
| --- | --- | --- |
| (GGS)_2_-mTagBFP2_Fwd | ACTAGTGGTGGTTCAGGAGGTAGTGTTTCAAAGGGTGAAGAG | #HH570 |
| mTagBFP2-ALFA-reverse | CATCTAGATGGTTCTGTTAAACGACGACGTAATTCTTCTTCTAATCTTGATGGTTTACTAGGTAAATCGCAGTAAC | #HH570 |
| NbALFA_fwd | AGGAGGTAGTAGATCTGAAGTACAATTACAAGAATCAGG | #HH574, #HH623, #HH686 |
| NbALFA_rev | TATTTATTTAACTAGTTGATGAAACAGTAACTTGTGTAC | #HH574, #HH623, HH676, #HH686 |
| pDM1501_HistoneH1_fwd | ATATAAAAAAAGATCTAAAAAATGGGTCCAAAAGCAC | #HH608 |
| pDM1501_mScarlet-I-ALFA_rev | TTTATTTTATACTAGTTGGTTCTGTTAAACGACGACGTAATTCTTCTTCTAATCTTGATGGCTTATACAATTCATCCATA | #HH608 |
| fbxD_fwd | AGGAGGTAGTAGATCTTCATATGACTATAACTGTTGGG | #HH687, #HH708 |
| fbxD_rev | ATTGTACTTCAGATCTTGGTACCATCTTAAATGTAC | #HH687, #HH708 |
| ecmAOp_fwd | ATTTAAAAAACTCGAGAATCATGGTAAAACAAATTG | #HH657 |
| ecmAOp_rev | CAAGATCTCAACGTTATAATTTTTAAAC | #HH657 |
| D19p_fwd | ATTTAAAAAACTCGAGGGTCCTAGAAAATTAAAAAA | #HH659 |
| D19p_rev | CCCATTTTTTAGATCTTAATTTATATTTACAAAAAATAATATAATATTTTGG | #HH659 |
| mAID_fwd | AATAAAAATCAGATCCAAAAAATGAAGGAAAAGTCTGCATG | #HH676 |
| mAID_rev | CATCTAGATTTATACATTCTCAAATC | #HH676 |
| mAID-NbALFA_fwd | GAATGTATAAATCTAGAGAAGTACAATTACAAGAATC | #HH676 |
| OsTIR(F74G)_fwd_1 | CAAGGATCCAAAAAATGACCTACTTCCCAG | #HH697 |
| OsTIR(F74G)_rev_1 | AATCGGCACCGTGAGGCTTACCCTTAAC | #HH697 |
| OsTIR(F74G)_fwd_2 | CTCACGGTGCCGATTTTAACTTGGTTC | #HH697 |
| OsTIR(F74G)_rev_2 | CATCTAGATAAAATCTTTACGAAGTTTGG | #HH697 |
| fpaAp_fwd | CAACTCGAGAGAATATGTGTCGCCCATAG | #HH350 |
| fpaA_rev | ACTAGTGTTTCCACCTTTATCTTCAC | #HH350 |
| fpaA_coding_fwd | CAAAGATCTAAAAAATGTCTTTAGTTAAATTAG | #HH698 |
| (GGS)_2_-OsTIR(F74G)_fwd | CATCTAGAGGTGGTTCAGGAGGTAGTACCTACTTCCCAGAAGAAG | #HH698 |
| OsTIR(F74G)_rev_3 | CTGAACCACCACTAGTTAAAATCTTTACGAAGTTTGG | #HH698 |
| pDM_NgoMIV_Foward | AAATGTCGAGGCCGGCCCACCCATGGTTTGAAAAT | #HH699, #HH700 |
| pDM_NgoMIV_Reverse | AACCGCCTCTGCCGGCATTTGAAAGGGCATCTGTTG | #HH699, #HH700 |
| pTM1285_acaA(4515-4534)_fwd | AGCATGCCTTCTTAACTTGAAAGA | #HH681 |
| pTM1285_acaA(4515-4534)_rev | AAACTCTTTCAAGTTAAGAAGGCA | #HH681 |
| acaA_fwd | AATAAAAATCAGATCTAAAAAATGGCATCTAGCTCACC | #HH722 |
| acaA_rev | CTGAACCACCACTAGTACTTGAAAGATGGAATCTTGG | #HH722 |
| acaA-homology-arm_(GGS)_2__fwd | TCTGGAACTCTCTTGGGTGAAATTGGCTCATTCACTACTCCAAGATTCCATCTTTCAAGTGGTGGTTCAGGAGGTAGT | – |
| acaA-homology-arm_ALFA-tag_rev | ATTTTTATTATTGACAATTGAATTTTTATAAATATATATTTGTTGTGTTTTGCCTTCTTATGGTTCTGTTAAACGAC | – |

### **Supplementary figures**

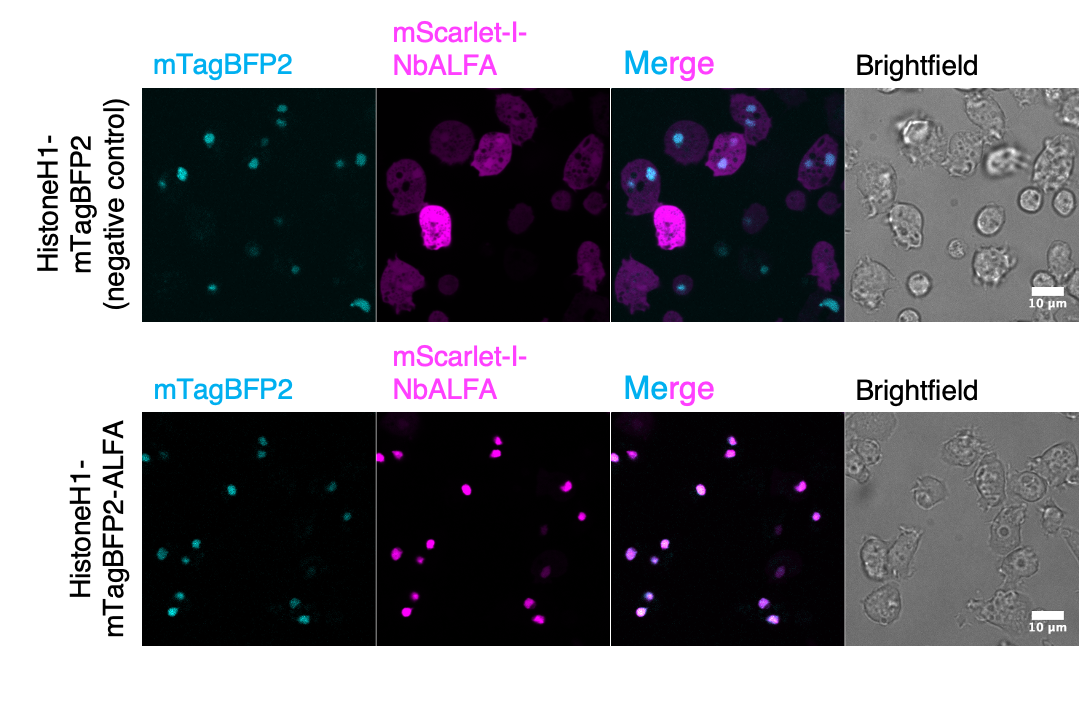

#### **Supplementary Fig. 1. Anti-ALFA nanobody (NbALFA) is sequestered to the nucleus by ALFA-tagged Histone H1.**

Snapshots of vegetative *D. discoideum* cells co-expressing mScarlet-I-NbALFA with either HistoneH1-mTagBFP2 (upper panels) or HistoneH1-mTagBFP2-ALFA (lower panels); (from left to right) fluorescence (cyan, Ex/Em = 405/447 nm; magenta, Ex/Em = 561/617 nm, merged) and brightfield images (grey scale). Scale bar, 10 µm. (B) Exogenous genes are expressed under the *act15* promoter.

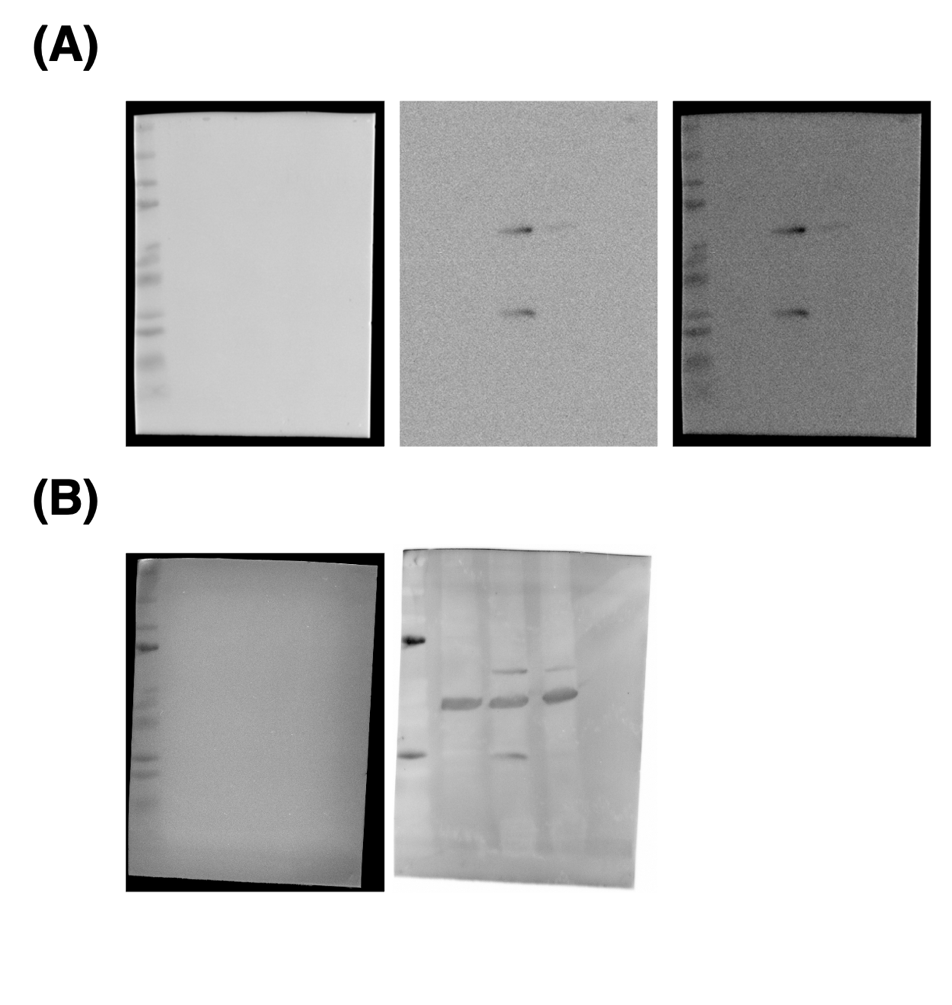

#### **Supplementary Fig. 2. Uncropped membrane western blot images related to Fig. 1.**

(A) Brightfield (left, marker), fluorescence (middle, anti-mouse-IgG goat antibody StarBrightBlue700) and merged images (right) obtained after incubation with an anti-RFP mouse monoclonal antibody. (B) Brightfield (left, marker) and fluorescence image (right, anti-mouse-IgG goat antibody StarBrightBlue700) acquired after (A), following subsequent incubation with an anti-actin mouse monoclonal antibody.

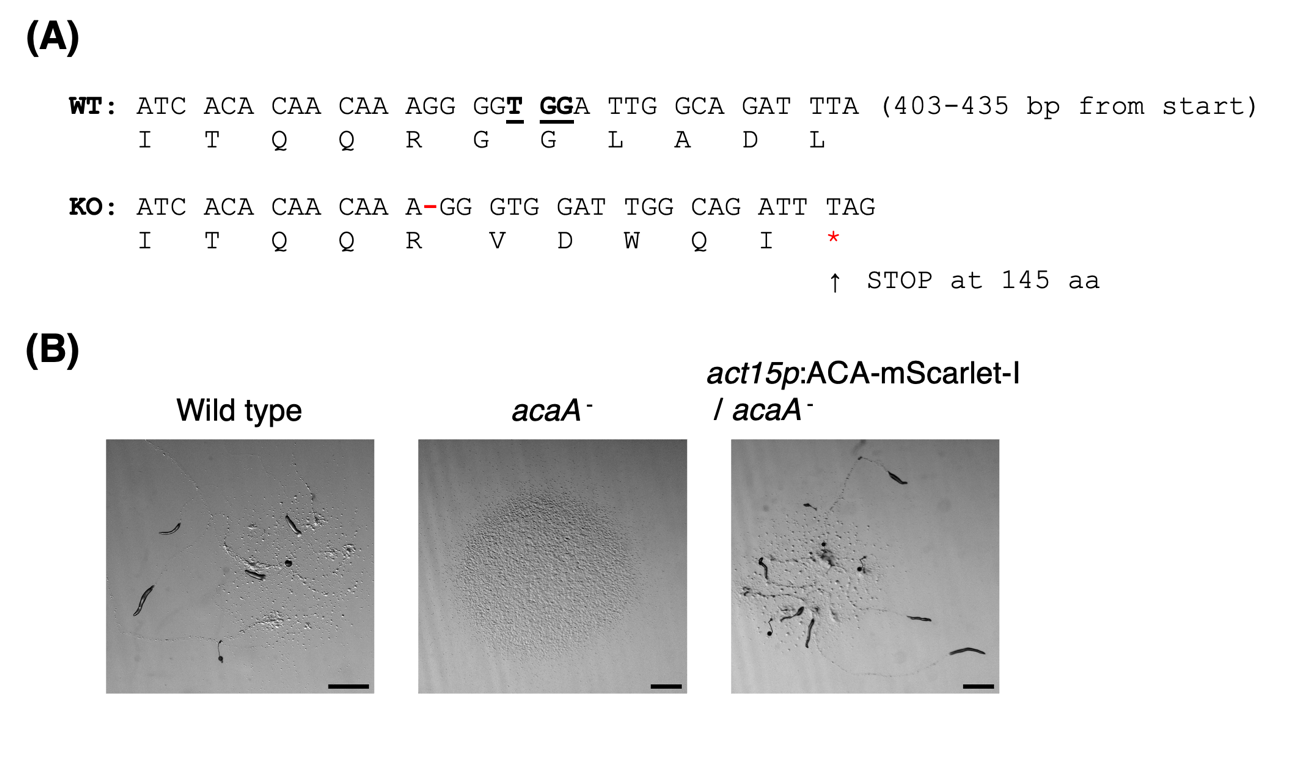

#### **Supplementary Fig. 3. Generation of an *acaA* knockout via CRISPR/Cas9.**

(A) DNA sequence of *acaA* in the parental Ax4 strain (WT) and an *acaA^-^* mutant (KO). PAM sequences are underlined. The mutant has a one-base deletion at position 416 (red hyphen) in the *ACA* gene, resulting in an amber codon at the 145th codon (red asterisk). (B) Snapshots of Ax4, *acaA^-^*, and the rescue strain expressing ACA-mScarlet-I under the *act15* promoter. Images were taken 24 hours after plating the cells on an agar plate. Scale bar, 1 mm.

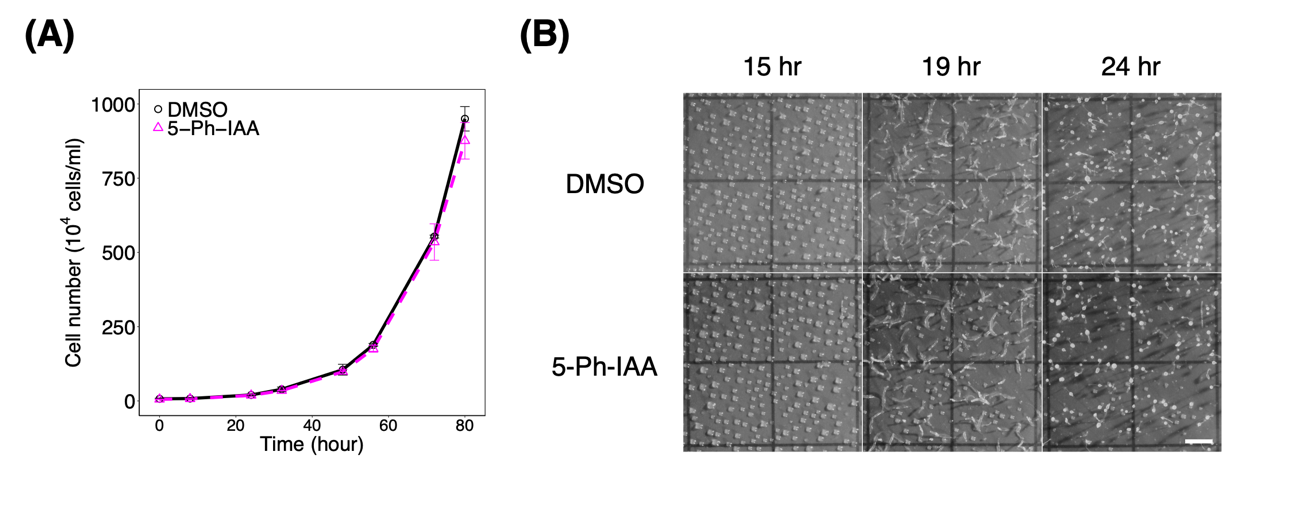

#### **Supplementary Fig. 4. 5-Ph-IAA does not interfere with the growth and development of *D. discoideum*.**

(A) Growth curves of Ax4 cells. 5-Ph-IAA solution in DMSO (magenta) or an identical volume of DMSO (mock control, black) was included in the growth medium. Bars indicate standard deviations. (B) Snapshots of developing Ax4 cells on a filter absorbed with buffer solution containing 5-Ph-IAA in DMSO or an equivalent volume of DMSO. Scale bar, 1 mm. Final concentration of 5-Ph-IAA was 10 µM for both (A) and (B).

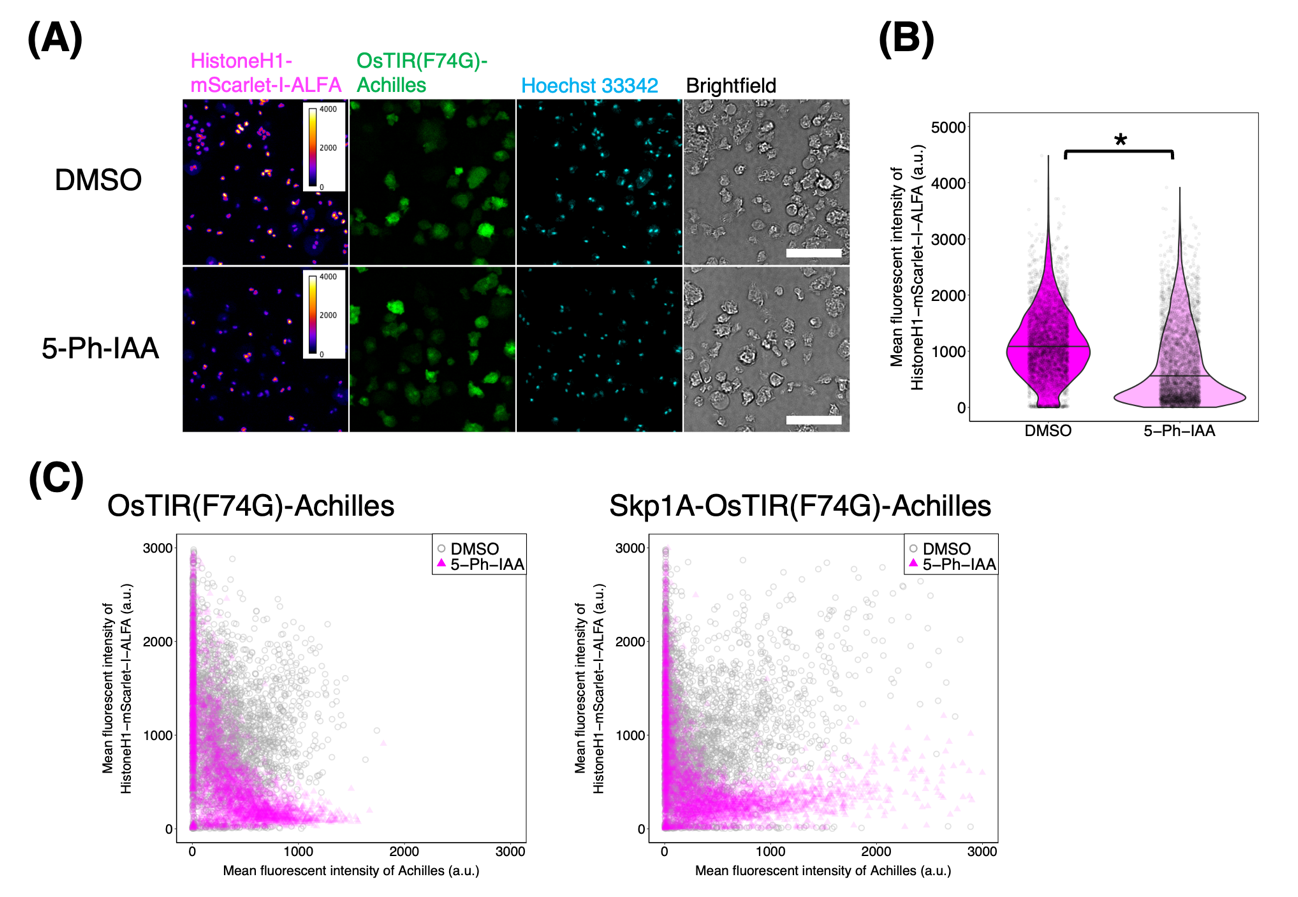

#### **Supplementary Fig. 5. OsTIR(F74G) without Skp1A fusion targets mAID-NbALFA-tagged Histone H1-mScarlet-I-ALFA in the presence of 5-Ph-IAA.**

(A) Snapshots of the *act5* locus in HistoneH1-mScarlet-I-ALFA knock-in cells carrying expression vectors *coaAp*:OsTIR(F74G)-Achilles and *act15p*:mAID-NbALFA. Vegetative cells were treated with DMSO (upper panels) or 10 µM 5-Ph-IAA (lower panels) for 24 hours, then fixed and labeled with Hoechst 33342. (Left to right) fluorescence images (heat color, Ex/Em = 561/ 617 nm; green, Ex/Em = 488/525 nm; cyan, Ex/Em = 405/447 nm) and brightfield (grey scale). Scale bar, 50 µm. (B) Violin plots of the mean fluorescence intensity of HistoneH1-mScarlet-I-ALFA in vegetative cells (DMSO, N = 3621 cells; 5-Ph-IAA, N = 3944 cells). The black line represents the median. **P* < 10^-15^. (C) The mean fluorescence intensity of HistoneH1-mScarlet-I-ALFA plotted against that of Achilles fused to OsTIR(F74G) or Skp1A-OsTIR(F74G) (OsTIR(F74G)–DMSO, N = 3540 nuclei. OsTIR(F74G)–5-Ph-IAA, N = 3803 nuclei. Skp1A–DMSO, N = 3712 nuclei. Skp1A–5-Ph-IAA, N = 3910 nuclei).

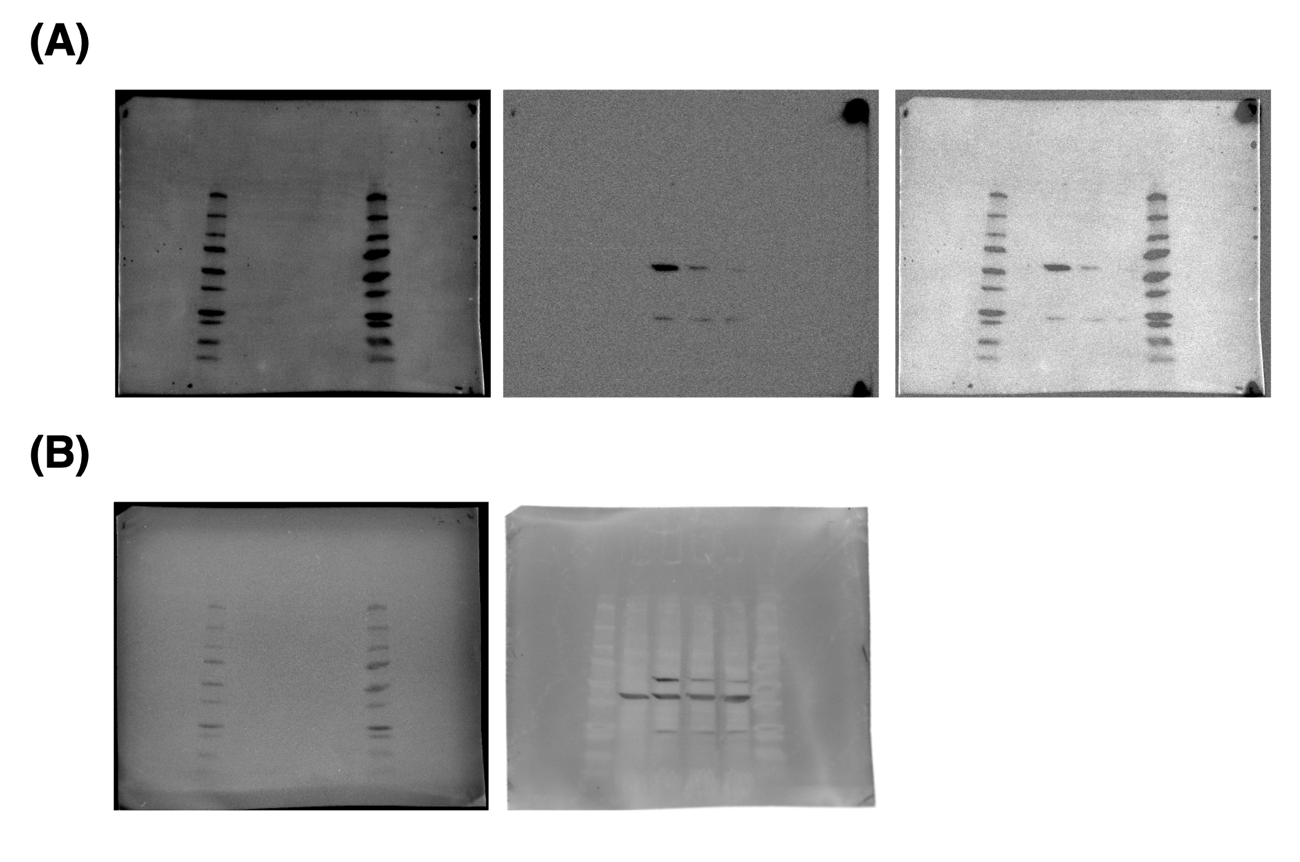

#### **Supplementary Fig. 6. Uncropped membrane western blot images related to Fig. 4.**

(A) Brightfield (left, marker), fluorescence (middle, anti-mouse-IgG goat antibody StarBrightBlue700) and merged images (right) obtained after incubation with an anti-RFP mouse monoclonal antibody. (B) Brightfield (left, marker) and fluorescence image (right, anti-mouse-IgG goat antibody StarBrightBlue700) acquired after (A), following subsequent incubation with an anti-actin mouse monoclonal antibody.

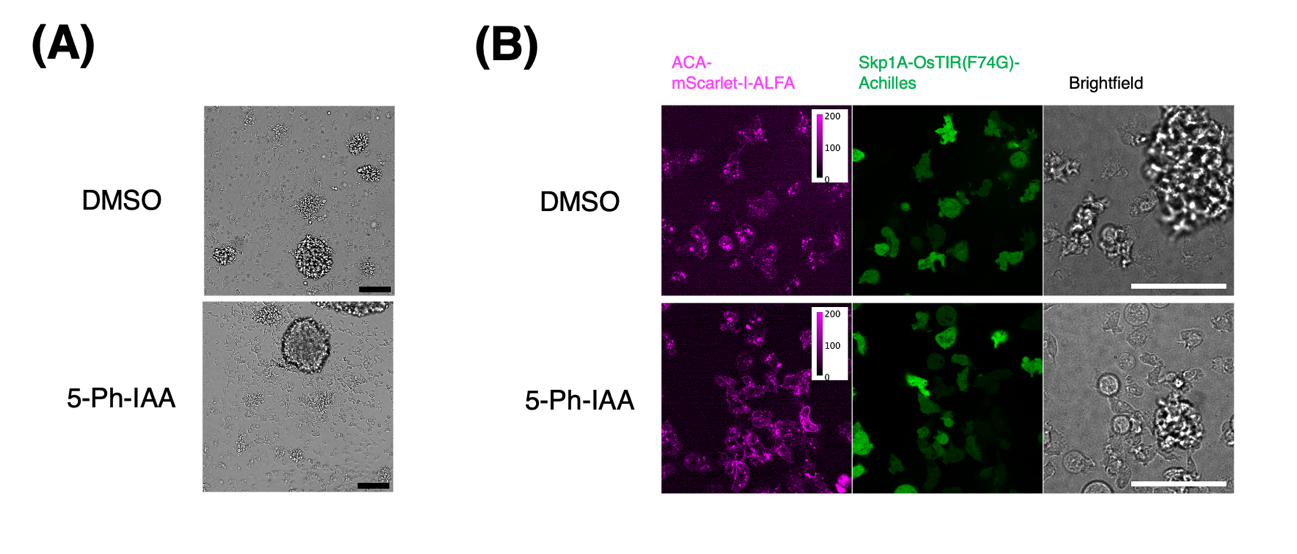

#### **Supplementary Fig. 7. Insufficient IAA-induced degradation of ACA-mScarlet-I-ALFA.**

ACA-mScarlet-I-ALFA knock-in cells carrying expression plasmids *coaAp*:Skp1A-OsTIR(F74G)-Achilles and *act15p*:mAID-NbALFA. (A) Low-magnification bright field images of cells in Developmental buffer treated with DMSO solution containing 5-Ph-IAA (lower panel; 10 µM final conc.) or DMSO only (upper panel) for 24 hours. Scale bar, 100 µm. (B) Fluorescence images (magenta, Ex/Em = 561/617 nm; green, Ex/Em = 488/525 nm) and brightfields of cells treated with DMSO (upper panel) or 10 µM 5-Ph-IAA (lower panel). Scale bar, 50 µm.
